# Unraveling the intricate microtubule inner protein networks that reinforce mammalian sperm flagella

**DOI:** 10.1101/2022.09.29.510157

**Authors:** Miguel Ricardo Leung, Marc C. Roelofs, Riccardo Zenezini Chiozzi, Johannes F. Hevler, Albert J. R. Heck, Tzviya Zeev-Ben-Mordehai

**Affiliations:** Structural Biochemistry Group, Bijvoet Centre for Biomolecular Research, Utrecht University, 3584 CG Utrecht, The Netherlands; Biomolecular Mass Spectrometry & Proteomics, Bijvoet Centre for Biomolecular Research and Utrecht Institute for Pharmaceutical Sciences, Utrecht University, 3584 CH Utrecht, The Netherlands

**Keywords:** sperm, motile cilia, microtubule inner proteins, cryo-electron microscopy

## Abstract

To find and fuse with the egg, mammalian sperm must complete an arduous voyage through the female reproductive tract. The sperm cell’s remarkable odyssey is powered by its flagellum, a microtubule-based molecular machine ornamented with accessory structures that stabilize the sperm tail in viscous media. Recently, cryo-electron tomography (cryo-ET) revealed that mammalian sperm flagella are further reinforced at the molecular scale with sperm-specific microtubule inner proteins (sperm-MIPs), but the identities of these sperm-MIPs are unknown. Here, we use cryo-electron microscopy to resolve structures of native bovine sperm doublet microtubules, thus identifying most sperm-MIPs. In the A-tubule, several copies of testis-specific Tektin-5 contribute to an extended protein network spanning nearly the entire microtubule lumen. Different copies of Tektin-5 adopt a range of conformations and organizations based on their local interactions with other MIPs. The B-tubule is in turn stabilized by sperm-MIPs that bind longitudinally along and laterally across protofilaments. We further resolve structures of endpiece singlet microtubules, revealing MIPs shared between singlets and doublets. Our structures shed light on the molecular diversity of cilia across different cell types of the vertebrate body and provide a structural framework for understanding the molecular underpinnings of male infertility.

## Main

To find and fuse with the egg, mammalian sperm must complete an arduous voyage through the female reproductive tract; along the way, they encounter viscous media, shear flows, and physical barriers ^1^. The sperm cell’s journey is powered by its flagellum, a microtubule-based molecular machine called a motile cilium. Motile cilia are microtubule-based assemblies used by a wide range of organisms and cell types to swim through fluid or to move fluid across their surfaces ^2,3^. The core of the motile cilium is the axoneme, a veritable molecular behemoth comprised of hundreds of different proteins, including dynein motors that drive motion and an extensive array of regulatory proteins that fine-tune motility ^4^. These proteins are anchored on axonemal microtubules, which consist of nine doublet microtubules (DMTs) arranged around a central pair of microtubules. High-resolution cryo-electron microscopy (cryo-EM) reconstructions of algal and mammalian DMTs revealed a panoply of microtubule inner proteins (MIPs) that stabilize the structure during ciliary beating ^5-10^. However, recent studies have begun to shed light on species- and cell type-specific differences in axoneme structure, particularly in the MIP repertoire ^11-14^.

Mammalian sperm flagella are longer (∼60 μm for bull sperm, ∼120 μm for mouse sperm; versus ∼7 μm for respiratory cilia, ∼10 μm for *Chlamydomonas* flagella) and stiffer than other ciliated cell types ^15,16^. This rigidity is due to the presence of large accessory structures like outer dense fibers and the fibrous sheath, which suppress buckling of the axoneme under loading in high-viscosity fluids like those in the female reproductive tract ^15-18^. Cryo-ET revealed that mammalian sperm axonemal DMTs and endpiece singlet microtubules (SMTs) are further reinforced at the nano-scale with extensive MIP densities ^12,19-21^, many of which remain unaccounted for in high-resolution structures of DMTs from *Chlamydomonas* flagella and mammalian respiratory cilia ^5,6^. It can be hypothesized that the additional MIPs in mammalian sperm (henceforth “sperm-MIPs”) serve to reinforce and thus stabilize the microtubule lattice itself against the large mechanical stresses involved in bending mammalian sperm flagella. However, because the identities of many sperm-MIPs are unknown, the structural mechanisms for such stabilization are unclear.

In this study, we use cryo-electron microscopy to resolve elaborate MIP networks in native mammalian sperm axonemal DMTs and endpiece SMTs. These reconstructions allow us to identify most sperm-MIPs and to describe their molecular organization, revealing how they reinforce the microtubule lattice. We further identify sperm-MIPs shared between axonemal DMTs and endpiece SMTs. Our structures can contribute towards identifying novel targets for contraception and will enhance our understanding of the molecular bases of male infertility.

### Cryo-EM reveals elaborate sperm-specific ornamentation of axonemal doublets

Mammalian sperm flagella present a challenge for purification and fractionation because axonemal microtubules are intimately associated with large accessory structures like the mitochondrial sheath, the outer dense fibers, and the fibrous sheath ^17^. Thus, rather than purifying sperm DMTs, we splayed the axonemes using a sliding disintegration protocol developed by Lindemann et al. ^22^. After the mitochondrial sheath is removed by DTT treatment and a freeze-thaw cycle, ATP-induced sliding causes the doublets to buckle out of the midpiece while still attached to their respective ODFs, disintegrating the axoneme and exposing individual DMTs suitable for cryo-EM **(Fig. 1a)**.

**Fig. 1.**
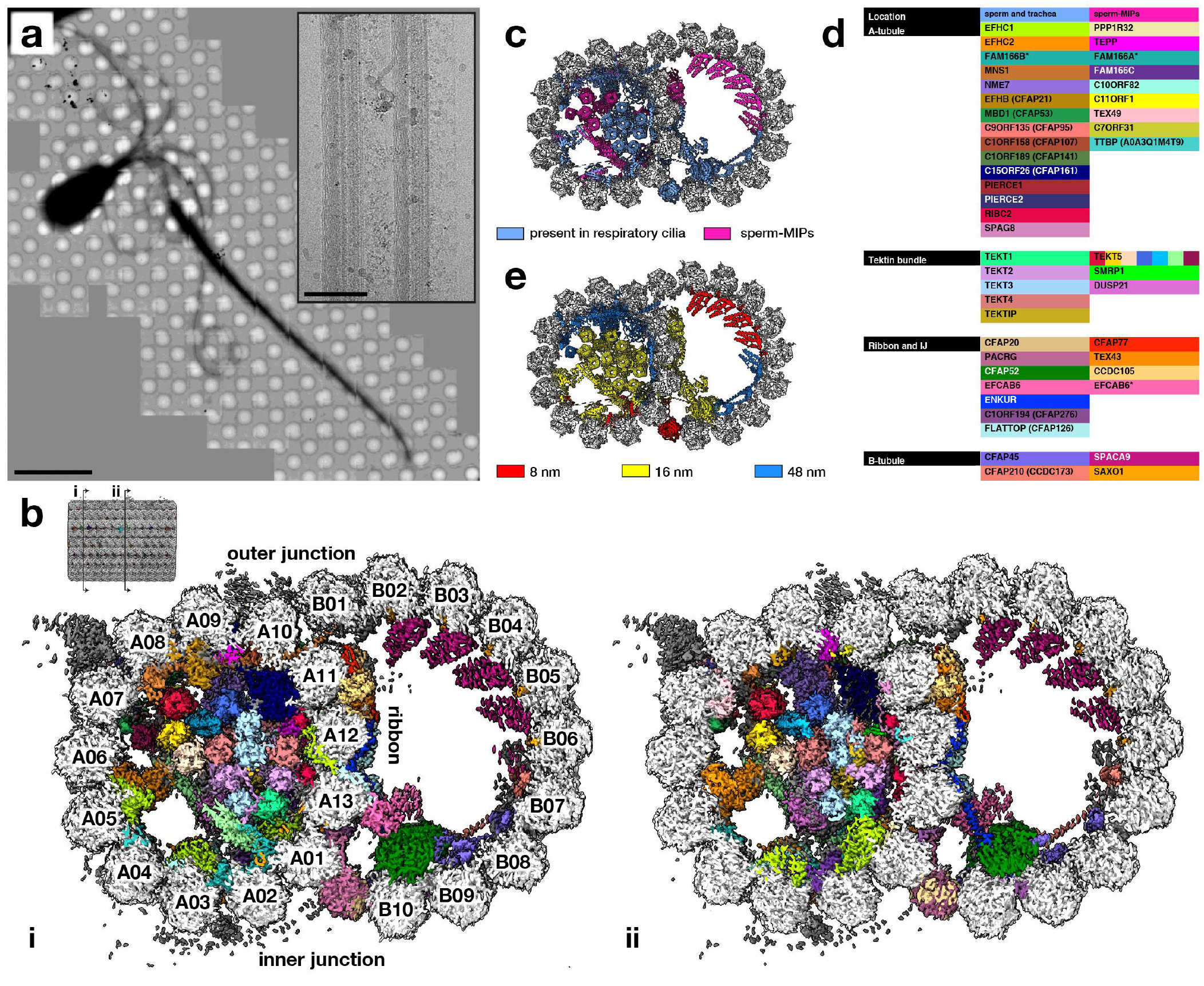
Cryo-electron microscopy unveils elaborate sperm-specific ornamentation of axonemal doublet microtubules (DMTs). **(a)** Cryo-EM image of a bovine sperm cell after sliding disintegration of the axoneme. Note how the DMTs are extruded from the midpiece. Scale bar: 10 μm. Inset: high-magnification cryo-EM image of sperm DMTs attached to their respective outer dense fibers. Scale bar: 100 nm. **(b)** Cryo-EM map of bovine sperm DMTs with 45 microtubule inner proteins (MIPs) colored individually; note that different groups of Tektin-5 are also colored individually (see Figs. 2 and 3). Panels (i) and (ii) are cross-sections taken at the locations indicated in the inset. **(c)** Atomic model of the 48-nm repeat of sperm DMTs with MIPs present in respiratory cilia colored in blue and sperm-MIPs colored in pink. **(d)** List of MIPs common to sperm and trachea (blue column) and MIPs newly-identified in sperm (pink column) arranged according to their locations in the DMT. **(e)** Atomic model of sperm DMTs with MIPs colored according to periodicity.

We reconstructed the 48-nm DMT repeat to an overall resolution of ∼3.7 Å, with local resolutions reaching ∼3 Å in the A-tubule lumen **(Fig. 1b, Fig. S1)**. Fitting our maps into *in situ* subtomogram averages of mammalian sperm DMTs showed that our structures retain all prominent MIP densities **(Fig. S2a)**, confirming that the sliding disintegration protocol preserves MIP architecture. We also resolve basal structures of external axonemal complexes like the radial spokes and the nexin/dynein regulatory complex **(Fig. S2b)**.

Based on well-resolved side chain density, we used the findMySequence program ^23^, supplemented with DALI searches and AlphaFold predictions, to confidently assign most MIP identities **(Fig. S3, Fig. S4)**, although a number of shorter or more poorly-resolved densities remain unassigned. Overall, our cryo-EM maps allowed us to build an atomic model consisting of >40 different MIPs **(Fig. 1b,d, Movie S1)**. Mammalian sperm DMTs retain nearly all MIPs present in respiratory cilia, one notable exception being FAM166B, which is expressed at low levels in the testis ^24^ and is replaced by FAM166A in sperm. Reconstructing the two halves of the 96-nm repeat confirmed the overall 48-nm periodicity of MIPs in sperm DMTs **(Fig. S2c)**. Individual sperm-MIPs have varying periodicities, but follow the general principles observed for bovine tracheal and *Chlamydomonas* DMTs, i.e. MIPs close to the A-tubule seam have 48-nm repeats and those in the A-tubule lumen have 16-nm repeats **(Fig. 1e)**.

At least 17 of the identified MIPs appear to be sperm-specific as they were not identified in previous cryo-EM maps of bovine respiratory cilia ^6^ **(Fig. 1c-d)**. We henceforth refer to the MIPs unique to sperm as “sperm-MIPs”. Sperm-MIPs form three major groups: MIPs found in the A-tubule **(Fig. 2, 3)**, MIPs at the ribbon (protofilaments A11-A13) and inner junction **(Fig. 7)**, and MIPs in the B-tubule **(Fig. 8)**, in addition to individual sperm-MIPs scattered across the DMT. This extensive ornamentation of the DMT may explain why we measure a more compact tubulin lattice in sperm DMTs versus tracheal DMTs **(Fig. S5)**.

**Fig. 2.**
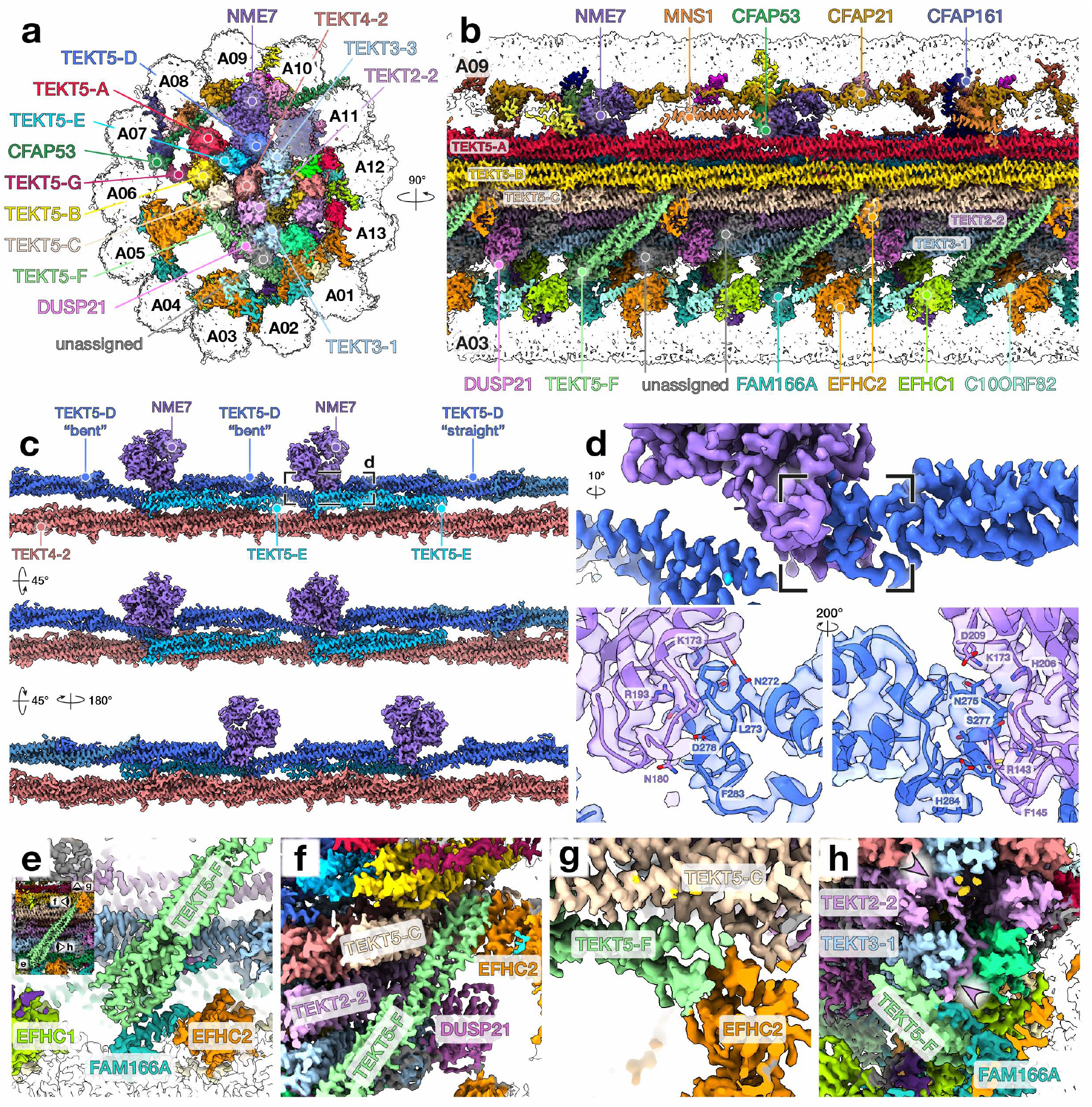
Multiple copies of the testis-specific Tektin-5 form an extended bundle in the A-tubule. **(a-b)** Cryo-EM map of the A-tubule of sperm DMTs with individual MIPs colored. **(c)** Interactions between NME7 and the TEKT5-D and TEKT5-E filaments. Note how the monomers in the TEKT5-D filament adopt different conformations depending on whether their 2A/2B helices interact with NME7 (“bent” TEKT5) or a neighboring tektin (“straight” TEKT5). **(d)** The L12 loop of the “straight” TEKT5-D monomer inserts into the putative nucleotide-binding pocket of NME7. The L12 of one “bent” TEKT5-D monomer interacts with the second copy of NME-7 in a similar fashion. Note that the top panel in (d) is rotated 10°relative to the view in (c). **(e-h)** Interactions between TEKT5-F and neighboring MIPs of the A-tubule. Viewing directions are indicated in the inset in panel (e).

**Fig. 3.**
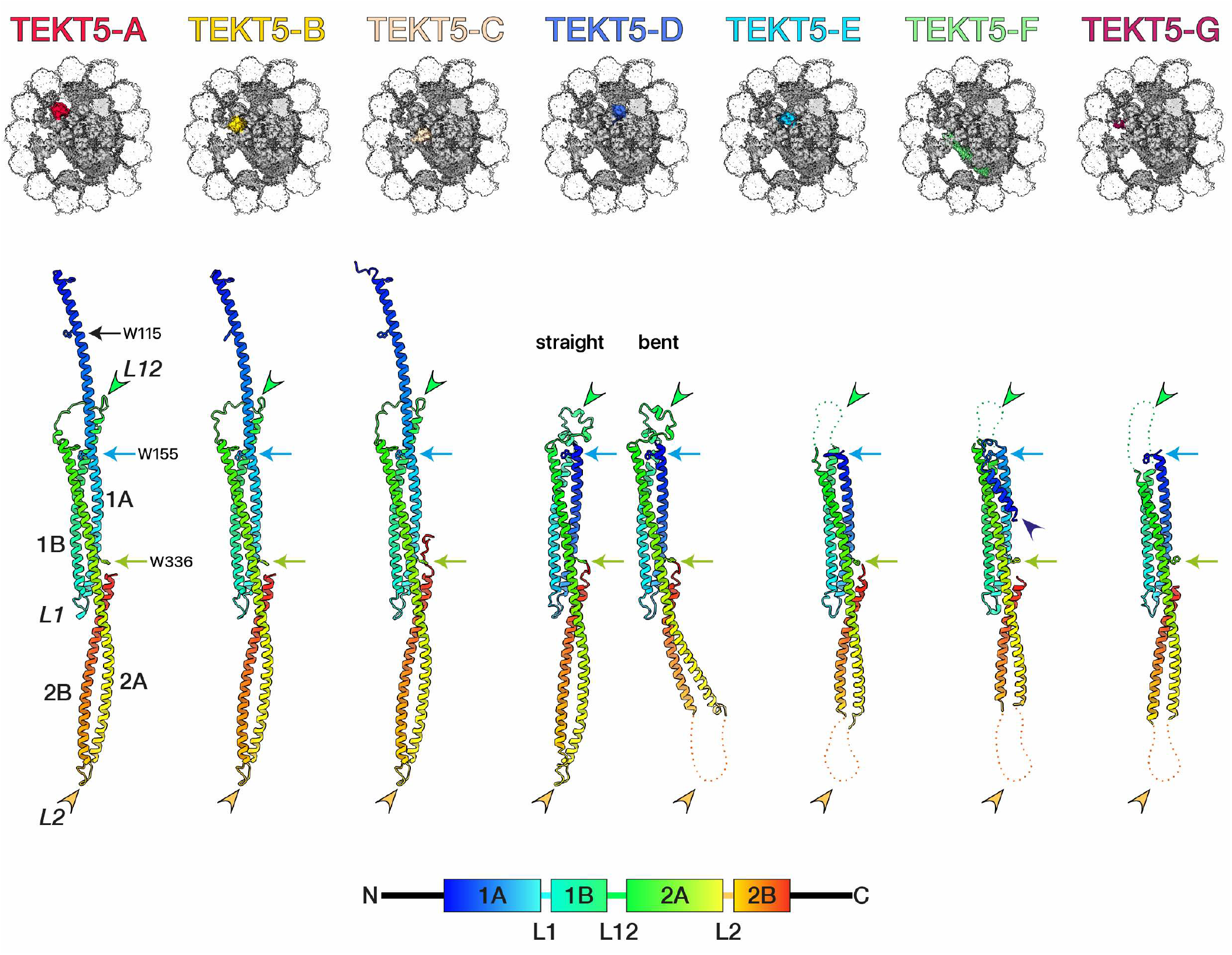
Multiple copies of Tektin-5 adopt different conformations based on their local interactions. For reference, the locations of Trp residues are indicated with black (Trp115, N-terminal region), blue (Trp155, 1A helix), and green (Trp336, 2A helix) arrows. A schematic of the tektin domain arrangement and corresponding color-code is provided at the bottom of the figure. The Tektin-5 filaments TEKT5-A, TEKT5-B, and TEKT5-C all adopt the canonical tektin conformation. Note how the N-terminal region is not resolved in the other groups, except in TEKT5-F where part of it refolds (dark blue arrowhead). In the group TEKT5-D, the L2 loop is disordered in the “bent” tektins, because these do not interact with other tektins; only the L2 loop of the “straight” tektin is resolved (compare orange arrowheads) because it is stabilized by interaction with a neighboring tektin (see also Fig. 2c). The L12 loops are ordered by interaction with the nucleotide-binding pocket of NME7 (compare green arrowheads, see also Fig. 2d).

### The sperm A-tubule has an extended tektin network formed by multiple copies of the testis-specific Tektin-5, which adopt a variety of conformations and organizations

Mammalian sperm have multiple unique alpha-helical MIPs in the A-tubule, which form an additional supercomplex (termed the “sickle” by Afzelius ^25^) attached to the 8-tektin bundle (the “pentagon”) **(Fig. 1c)**. As a result, nearly the entire lumen of the A-tubule is filled with MIPs **(Fig. 1b, 2a)**, explaining its dense appearance in electron micrographs. Well-resolved side chain density allows findMySequence ^23^ to confidently assign the alpha-helical MIPs as multiple copies of the testis-specific Tektin-5 (TEKT5). These copies form seven groups (TEKT5-A to TEKT5-G) based on their location and organization. Three groups of Tektin-5 (TEKT5-A: red, TEKT5-B: yellow, TEKT5-C: beige) form filaments through the canonical end-to-end association seen in other tektins ^6^ (i.e. the L2 loop of one molecule inserts into the L12 loop of the adjacent molecule) **(Fig. 2b, 4)**.

**Fig. 4.**
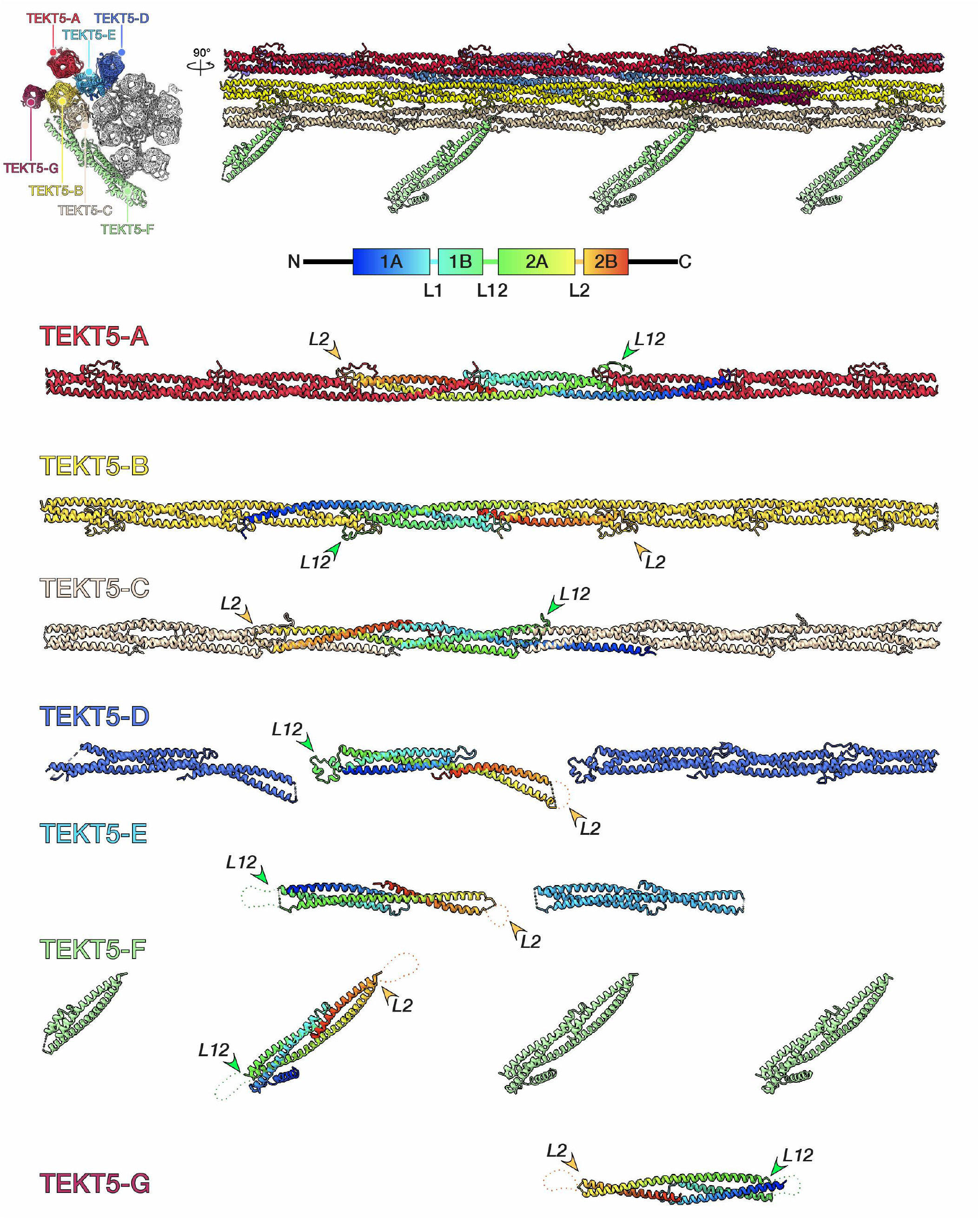
Comparing the quaternary structures of Tektin-5 in the bovine sperm DMT. Tektins are grouped according to similar positions in the DMT, with groups labelled from “A” to “G”. For each group, one monomer is colored in a rainbow palette from N-(blue) to C-terminus (red).

The other four groups of Tektin-5 (TEKT5-D to TEKT5-G) adopt novel conformations and organizations. For example, TEKT5-D (dark blue) near NME-7 forms an interrupted filament; in every 48-nm repeat, one copy associates with its neighbor via the canonical interface (“straight” tektin), but two copies adopt a “bent” conformation, where their 2A/2B helices curve downwards to accommodate NME-7 **(Fig. 2c, 4)**. These bent helices are then positioned to interact with TEKT4-2 in the pentagon **(Fig. 2c)**. One “bent” Tektin-5 molecule and one “straight” Tektin-5 molecule each interact directly with one copy of NME-7, their L12 loops inserting into the NME-7 putative nucleotide-binding pocket **(Fig. 2d)**. Two additional “bent” Tektin-5 molecules (TEKT5-E: light blue) found near the two copies of NME-7 bridge TEKT5-A, TEKT5-B, TEKT5-D, and TEKT4-2 **(Fig. 2a,c)**.

Four copies of Tektin-5, TEKT5-F, are unique in that they do not run parallel to the other tektin filaments. Instead, they run diagonally perpendicular to the long axis of the DMT, with one copy every 16-nm forming the handle of the “sickle” **(Fig. 2a-b, 4)**. Their N-termini interact with FAM166A between protofilaments A01-A02 **(Fig. 2e)**, and their C-termini interact with a groove formed by an interaction between the N-terminus of TEKT5-C and EFHC2 **(Fig. 2f,g)**. The C-terminal tail of TEKT2-2 also loops around TEKT3-1 to interact with TEKT5-F **(Fig. 2h)**. In addition, three densities bind to TEKT2-2 and TEKT3-1 near TEKT5-F **(Fig. 2b)**. By building a partial poly-alanine model and querying the DALI server, we could assign one of these densities as a dual-specificity phosphatase domain, which findMySequence subsequently identified as DUSP21 **(Fig. 2b, S3c)**. The two densities flanking DUSP-21 remain unassigned. Finally, in every 48-nm repeat there is one additional copy of Tektin-5, TEKT5-G, found between TEKT5-B and CFAP53 close to protofilament A06 **(Fig. 2a, 4)**. TEKT5-G density is weaker than adjacent Tektin-5 molecules, suggesting that it may only be weakly bound to its neighbours or that it is only present in a subset of DMTs.

Comparing the models of the Tektin-5 filaments derived from our cryo-EM map reveals unexpected diversity in tektin conformation **(Fig. 3)**. As described above, individual molecules within the TEKT5-A, TEKT5-B, and TEKT5-C filaments interact through the canonical mechanism seen in Tektin-1 to -4. As such, Tektin-5 monomers within these filaments adopt the conformations described previously. However, the monomers within the TEKT5-D, TEKT5-E, TEKT5-F, and TEKT5-G groups do not directly interact with one another, so they adopt different conformations. For instance, the L12 loop – which normally wraps around the L2 loop of the neighboring molecule – is disordered in TEKT5-E, TEKT5-F, and TEKT5-G **(Fig. 3, green arrowheads)**. It is ordered in the TEKT5-D group because it is stabilized by interactions with NME-7 **(Fig. 2c and Fig. 3, green arrowheads)**. Likewise, the L2 loops are disordered in most of these groups **(Fig. 3, orange arrowheads)**, except for the “straight” protomer in TEKT5-D, where the L2 loop is ordered because it is stabilized by interactions with a neighboring Tektin-5 **(Fig. 2c)**. Likewise, the Tektin-5 molecules that do not interact directly with other tektins lack density before the region of Trp155 **(Fig. 3, blue arrows)**. This corresponds to ∼60 fewer residues resolved when compared to the Tektin-5 molecules that form filaments, where the N-terminal region is stabilized by packing against the adjacent molecule. One exception is in TEKT5-F, which binds diagonally across the A-tubule, where we resolve ∼30 more residues before Trp 155 **(Fig. 3, dark blue arrowhead)**. In TEKT5-F, this region of the 1A helix has refolded into a helix and a loop, forming a hairpin which interacts with FAM166A **(Fig. 2e)**.

### Sperm-MIPs interact with the tektin bundle, forming a continuous protein interaction network that bridges the A-tubule lumen

In addition to the Tektin-5 bundle, multiple individual sperm-MIPs are scattered throughout the A-tubule **(Fig. 1b-d, 5, 6)**. For example, C11ORF1 and TEPP (testis-, prostate-, and placenta-expressed) are novel seam-binding MIPs bridging protofilaments A09 and A10 **(Fig. 5a)**. TEX49 binds at protofilaments A07/A08 close to Pierce1/2; although structurally-unrelated to the Pierce proteins, TEX49 also extends out of the DMT and appears to interact with coiled-coil proteins that form the base of the outer dynein arm docking complex **(Fig. 5b)**.

**Fig. 5.**
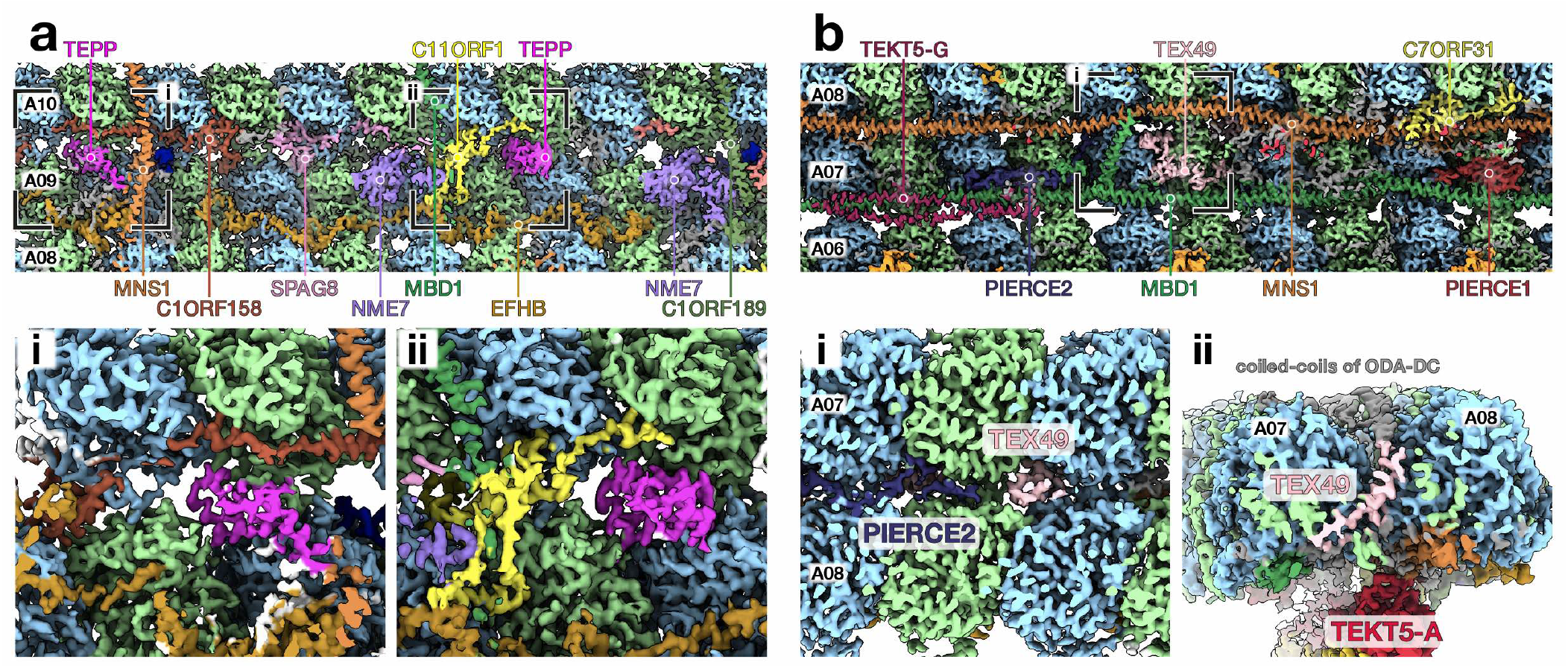
Sperm-MIPs in the A-tubule and their interactions with the tubulin lattice. **(a)** MIPs bound to protofilaments A09/A10, the location of the A-tubule seam. The sperm-MIPs C11ORF1 (yellow) and TEPP (testis-, prostate-, and placenta-expressed, bright pink) are novel seam-binding MIPs. **(b)** MIPs bound to protofilaments A07/A08, whose external inter-protofilament ridge serves as the binding site for the outer dynein arm docking complex (ODA-DC). The sperm-MIP TEX49 (testis-expressed 49) extends through the microtubule wall and appears to interact with the coiled-coils at the base of the ODA-DC.

Around protofilaments A01/A02, FAM166C docks onto TEKT1-1 with a 16-nm repeat **(Fig. 6a-b)**. A helix close to the FAM166C N-terminus packs against the 1B and 2A helices of TEKT1-1 **(Fig. 6c)**, while the FAM166C C-terminus extends downwards to interact with EFHC1 **(Fig. 6a)**. Also interacting with TEKT1-1 is PPP1R32 (protein phosphatase 1 regulatory subunit 32), which interacts with the TEKT1-1 L1/2 loop at the inter-protomer interface once every 48 nm **(Fig. 6d)**. PPP1R32 bridges TEKT1-1 and the tubulin lattice, interacting with both the inter- and intra-dimer α/β-tubulin interfaces **(Fig. 6e)**. Notably, PPP1R32 was identified as a conserved ciliary protein in an evolutionary proteomics study, where it was also confirmed to localize to the ciliary base when heterologously expressed in cultured cells ^26^.

**Fig. 6.**
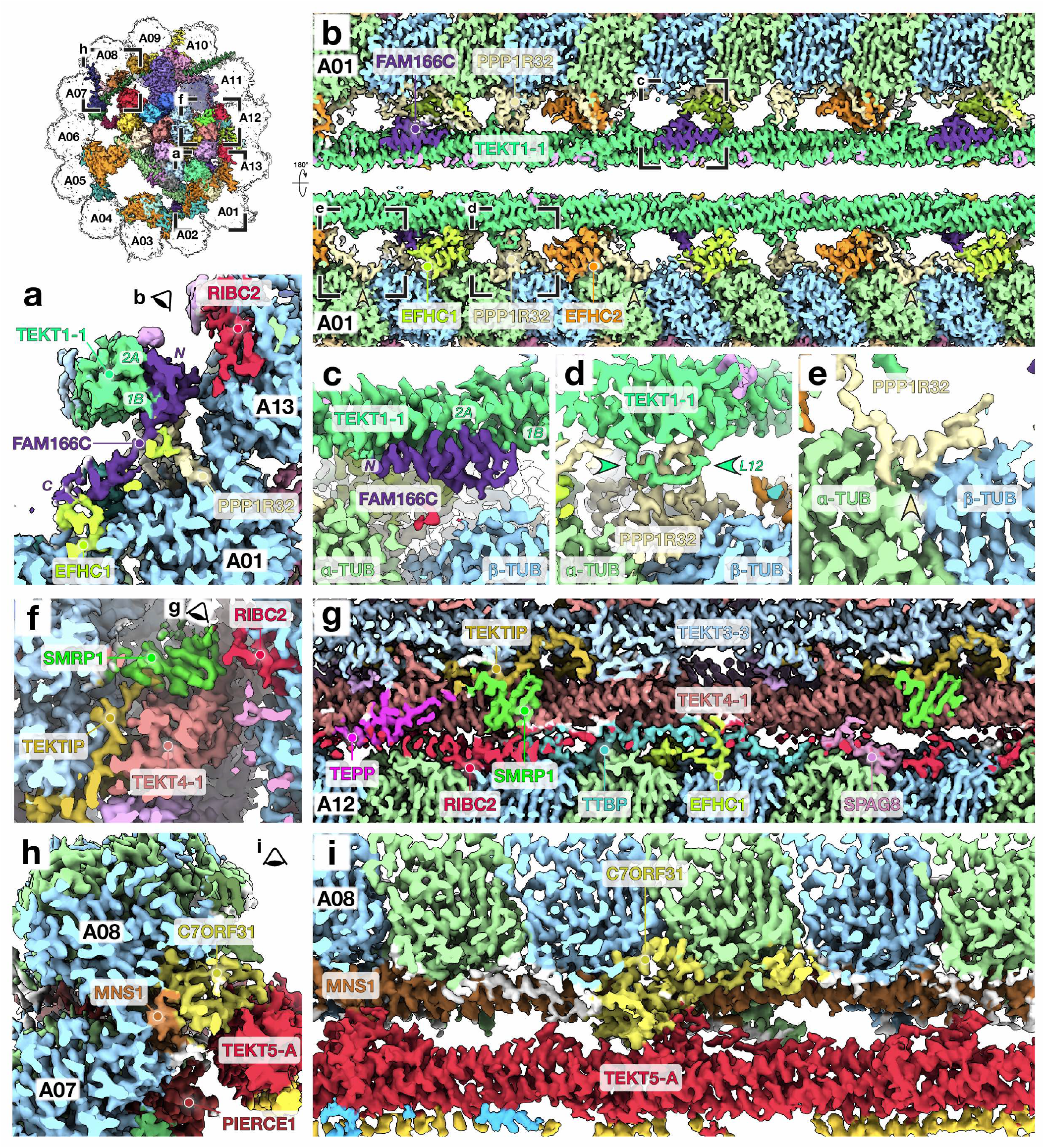
Sperm-MIPs in the A-tubule bridge the tektin bundle and the tubulin lattice, forming an interconnected protein network that spans nearly the entire microtubule lumen. **(a)** At protofilament A01, Tektin1-1 (sea green) interacts with two sperm-MIPs: FAM166C (family with sequence similarity 166 member C, purple) and PPP1R32 (protein phosphatase 1 regulatory subunit 32, pale yellow). FAM166C bridges Tektin1-1 and EFHC1 on protofilament A02, while PPP1R32 bridges Tektin1-1 and the tubulin lattice at protofilament A01. **(b)** FAM166C repeats once every 16-nm while PPP1R32 repeats once every 48-nm. **(c-e)** Close-up views of interactions between Tektin1-1 helices 1B/2A and FAM166C (c); between the Tektin1-1 L12 loop and PPP1R32 (d); and between the intra-dimer α-/β-tubulin interface and PPP1R32 (e). **(f)** Beside protofilament A12, the sperm-MIP SMRP1 (spermatid-specific manchette-related protein 1) packs against Tektin4-1 and interacts with the N-terminus of TEKTIP. **(g)** SMRP1 has a 16-nm repeat. Other sperm-MIPs in this region, TEPP and TTBP (tektin- and tubulin-bridging protein, UniProt A0A3Q1M4T9) have 48-nm repeats. TTBP bridges Tektin4-1 and the tubulin lattice at protofilament A12. **(h-i)** The sperm-MIP C7ORF31 bridges Tektin5-A and the tubulin lattice at protofilament A08.

Beside protofilament A12, the sperm-MIP SMRP1 (spermatid-specific manchette-related protein 1) interacts with TEKT4-1 and with the N-terminus of TEKTIP **(Fig. 6f)**. SMRP1 repeats every 16-nm, but every 48-nm there is one copy of TEPP that rests on top of TEKT4-1 close to the SMRP1 binding site **(Fig. 6g)**. Another MIP with 48-nm periodicity binds in this region -the uncharacterized protein UniProt A0A3Q1M4T9, which bridges TEKT4-1 and tubulin in protofilament A12 **(Fig. 6g)**. We therefore rename this protein the “tubulin-and tektin-bridging protein” or TTBP. We note that proteomics identified both TTBP and PPP1R32 in bovine respiratory cilia ^6^; however, the proteins were not identified in the corresponding cryo-EM maps, possibly because they have a more limited expression pattern around or along the axoneme in tracheal cilia as opposed to sperm.

On the opposite side of the A-tubule, the sperm-MIP C7ORF31 binds to tubulin in protofilament A08, as well as to MNS1 and TEKT5-A **(Fig. 6h-i)**. C7ORF31 may also function in the centriole, as it was identified in a proteomics study of sperm centrioles and shown to localize to centrosomes when expressed in cultured cells ^27^. Altogether, with C7ORF31 interacting with TEKT5-A, PPP1R32 interacting with TEKT1-1, EFHC2 interacting with TEKT5-C and TEKT5-F, and NME-7 interacting with TEKT5-D, the sperm-MIPs in the A-tubule form an interconnected protein network bridging nearly the entire microtubule lumen.

### Sperm-MIPs at the ribbon and inner junction interact with tubulin C-terminal tails

The inner junction (IJ), where A01 and B10 are connected by alternating copies of FAP20 and PACRG, is highly conserved between algae and mammals ^5,6,8^. The only major addition identified in bovine respiratory cilia was one copy of EFCAB6 bound to CFAP52 every 48-nm ^6^. However, in sperm, the IJ and its neighboring protofilaments A11-A13 – known as the ribbon – are supplemented with additional MIPs **(Fig. 1c, Fig. 7)**. In sperm, there are two additional copies of EFCAB6, which thus has a 16-nm repeat as opposed to the 48-nm repeat in respiratory cilia **(Fig. 7a)**. There is also a prominent filament of alpha-helical MIPs bound to the inter-protofilament ridges between A11 and A12, which we identify as the protein CCDC105. Two additional sperm-MIPs bridge the IJ with CCDC105: CFAP77 binds between FLATTOP and CCDC105; TEX43 binds close to ENKUR and wraps around the inter-protomer interface of CCDC105. We resolve additional density for the C-terminal tails (CTTs) of the tubulin molecules to which these sperm-MIPs are bound **(Fig. 3b, arrowheads)**. The CTT of α-tubulin packs against TEX43, while the CTT of β-tubulin from the neighboring dimer is sandwiched between CFAP77 and ENKUR.

**Fig. 7.**
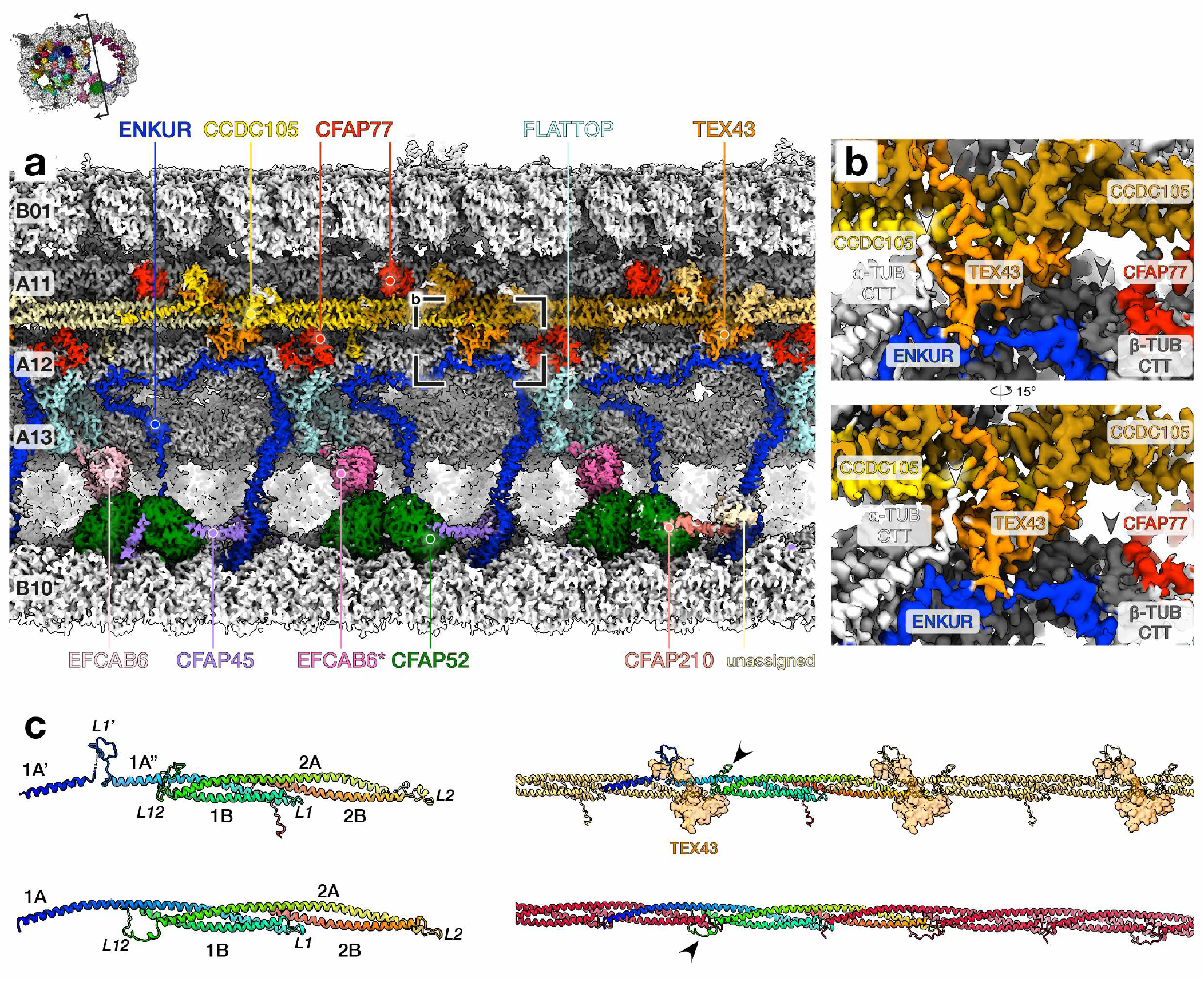
Sperm-MIPs at the ribbon interact with tubulin C-terminal tails. **(a)** Cryo-EM map of the ribbon and inner junction of sperm DMTs with MIPs colored individually. Each protomer in the CCDC105 filament is colored separately for clarity. **(b)** Additional densities attributable to the C-terminal tails (CTTs) of tubulin are clearly visible on protofilament A12, near sperm-MIPs TEX43, CFAP77, and CCDC105. TEX43 and CFAP77 appear to interact with the CTTs of alpha- and beta-tubulin respectively (white and grey arrowheads). **(c)** Comparing the tertiary (left) and quaternary (right) structures of CCDC105 (upper) and tektin filaments (lower, tektin 5-A as an example). In the left panels, proteins are colored in a rainbow palette from N- (blue) to C-terminus (red). CCDC105 secondary structure is annotated by analogy to tektin. In the right panel, one copy of each protein is colored in a rainbow palette from N to C. The arrowheads indicate differences in the inter-protomer interface. For the CCDC105 filament, TEX43 molecules that bind at the inter-protomer interfaces are shown as transparent surfaces.

The tertiary structure of CCDC105 is similar to that of the tektins **(Fig. 7c)**. However, instead of having four helices, CCDC105 has five – what would be the equivalent of the tektin 1A helix is instead split into two helices (which we call 1A’ and 1A”) separated by a loop (L1’), with the N-terminal 1A’ helix being shorter **(Fig. 7c, upper left panel)**. Furthermore, the mechanism by which protomers assemble into filaments differs slightly between tektins and CCDC105 **(Fig. 7c, right)**. Like the tektins, the main points of contact between neighboring CCDC105 molecules are helical overhangs that extend beyond the central bundle. However, CCDC105’s equivalent of the L12 loop does not clamp around the L2 loop of the neighboring molecule, instead looping around the 1A” helix of the same protomer **(Fig. 7c, arrowheads)**. Interactions between neighboring CCDC105 molecules may be further stabilized by the sperm-MIP TEX43, which acts like a staple, looping around the inter-CCDC105 interface and interacting with protofilaments A11 and A12 **(Fig. 7a-c)**. Functionally, little is known about CCDC105 other than that it is highly-expressed in testis ^24,28^ and that it is down-regulated in a mouse QRICH2-knockout model exhibiting multiple morphological abnormalities of the flagella ^29^.

### Sperm-MIPs in the B-tubule longitudinally and laterally reinforce the microtubule lattice

Another major addition to sperm DMTs are prominent MIPs bound to protofilaments B02-B07 of the B-tubule **(Fig. 8a)**, which lack large MIPs in DMTs from either bovine trachea or *Chlamydomonas*. These B-tubule MIPs correspond to the ladder-like densities resolved in *in situ* subtomogram averages of pig **(Fig. 9a)** and horse sperm DMTs **(Fig. 9b)**. One B-tubule MIP (orange) presents as a filamentous density binding at the intradimer interface with an apparent ∼8-nm periodicity. The binding site, repeat distance, and morphometry of this density closely resemble known MIPs FAP363 from *Chlamydomonas* ^5^ and SPM1 from *Toxoplasma gondii* ^30^, which are members of the MAP6-related family of microtubule-stabilizing proteins **(Fig. 10)** ^31,32^. Specifically, the density is consistent with the conserved Mn motif present in this family – a short helix flanked by a conserved tyrosine/phenylalanine on one end and a threonine/serine on the other **(Fig. 10a)** ^30,31^.

**Fig. 8.**
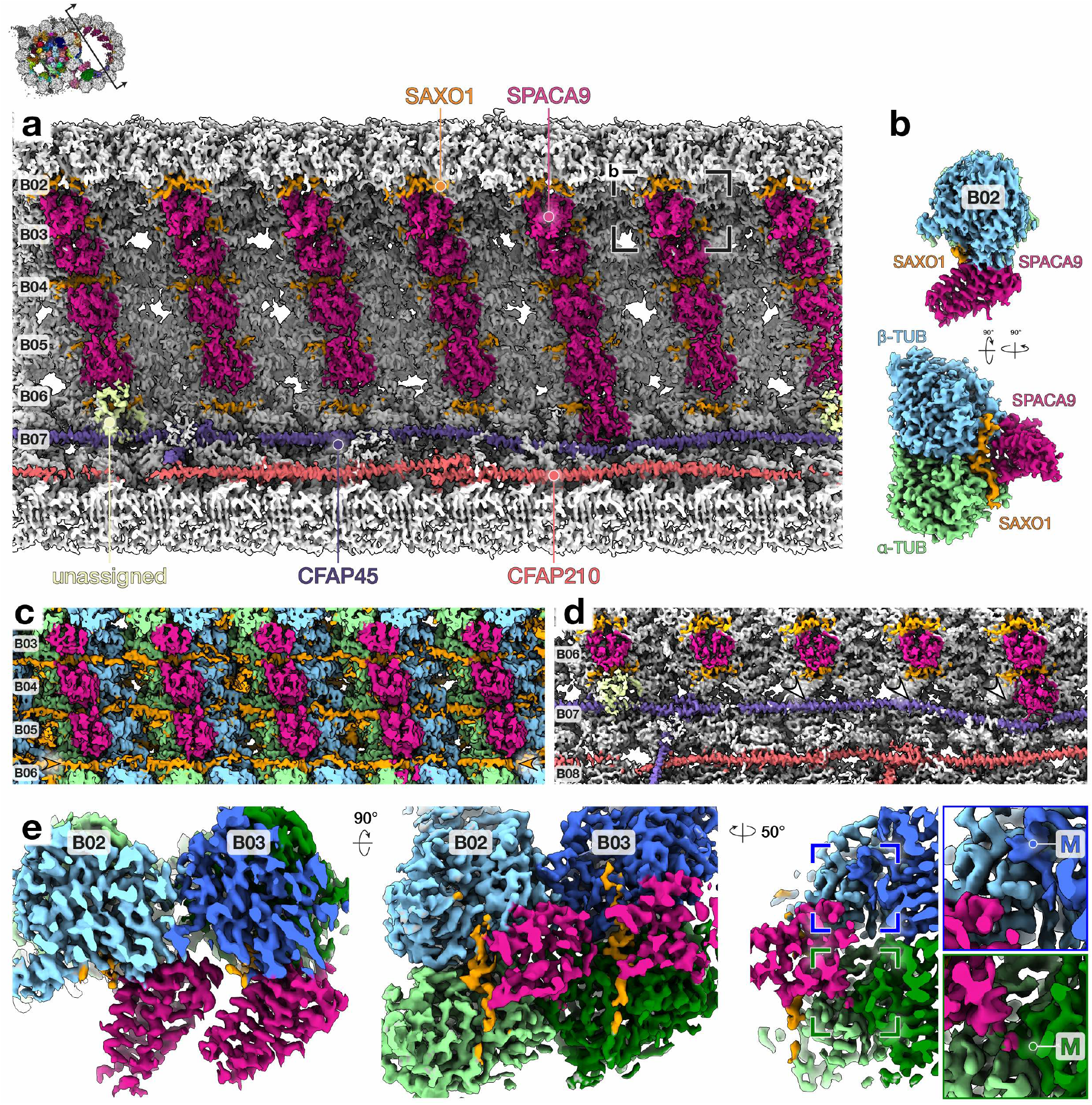
The B-tubule is decorated with MIPs that interact longitudinally and laterally across protofilaments. **(a)** Cryo-EM map of the B-tubule of sperm DMTs, with individual MIPs colored. **(b)** Extracted density for one tubulin dimer and associated MIPs from protofilament B02, specifically from the region indicated in (a). **(c)** Unsharpened cryo-EM map of protofilaments B03-B06 illustrating how SAXO1 appears to bind longitudinally along individual protofilaments.**(d)** Every 48 nm, an additional copy of SPACA9 is present at the B06/B07 interface, where CFAP45 bends away and exposes a binding site (compare arrowheads). The unassigned MIP binds even when CFAP45 does not bend away, suggesting it may recognize CFAP45 instead of the B06/B07 interface. **(e)** Extracted density for neighboring tubulin dimers and associated SAXO1 and SPACA9 molecules taken from protofilaments B02/B03. SPACA9 interacts with both α- and β-tubulin, along with the M-loop (labelled “M”) of α-tubulin from the neighboring protofilament (green box) but not the M-loop of the neighboring β-tubulin (blue box).

**Fig. 9.**
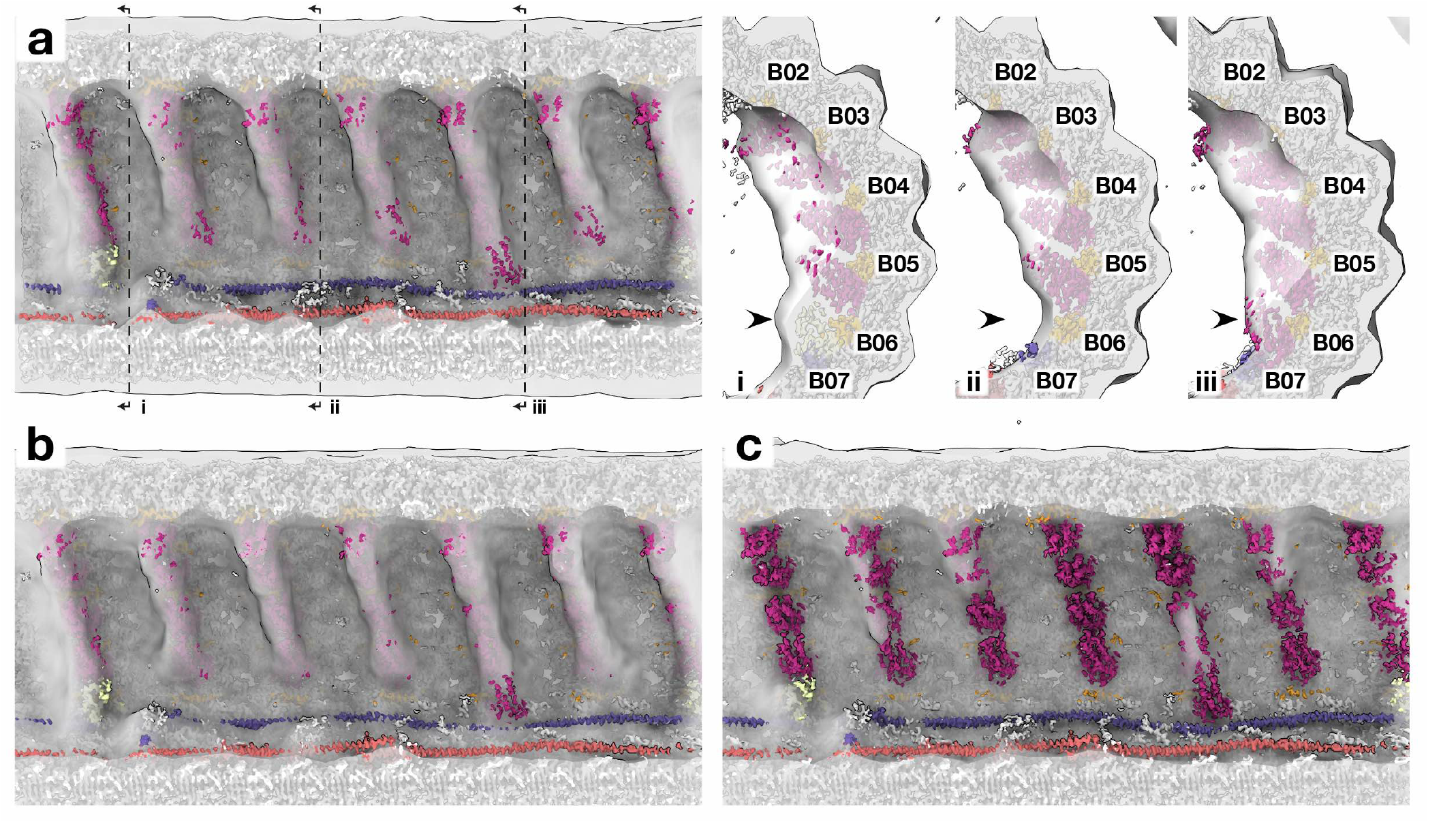
Prominent ladder-like densities in the B-tubule are present in pig sperm and horse sperm, with shorter shelf-like protrusions mouse sperm. Single-particle cryo-EM reconstruction of bovine sperm DMTs (this study) docked into in situ subtomogram averages of **(a)** pig sperm (EMD-12070), **(b)** horse sperm (EMD-12078), and **(c)** mouse sperm (EMD-12133). In (a), panels (i) to (iii) illustrate how the pattern of long and short ladder rungs visible in the subtomogram averages is consistent with the periodicity observed in the high-resolution single-particle structure. The ladder rungs are formed by neighboring copies of SPACA9, which bind close to the inter-protofilament interface. In every 48-nm repeat, there are 6 ladder rungs. There is one long rung (four copies of SPACA9 from B02-B06 plus one unassigned protein at B06/B07) (i), followed by three short rungs (four copies of SPACA9 from B02-B06) (ii), followed by another long rung (five copies of SPACA9 from B02-B07) (iii), followed finally by one short rung.

**Fig. 10.**
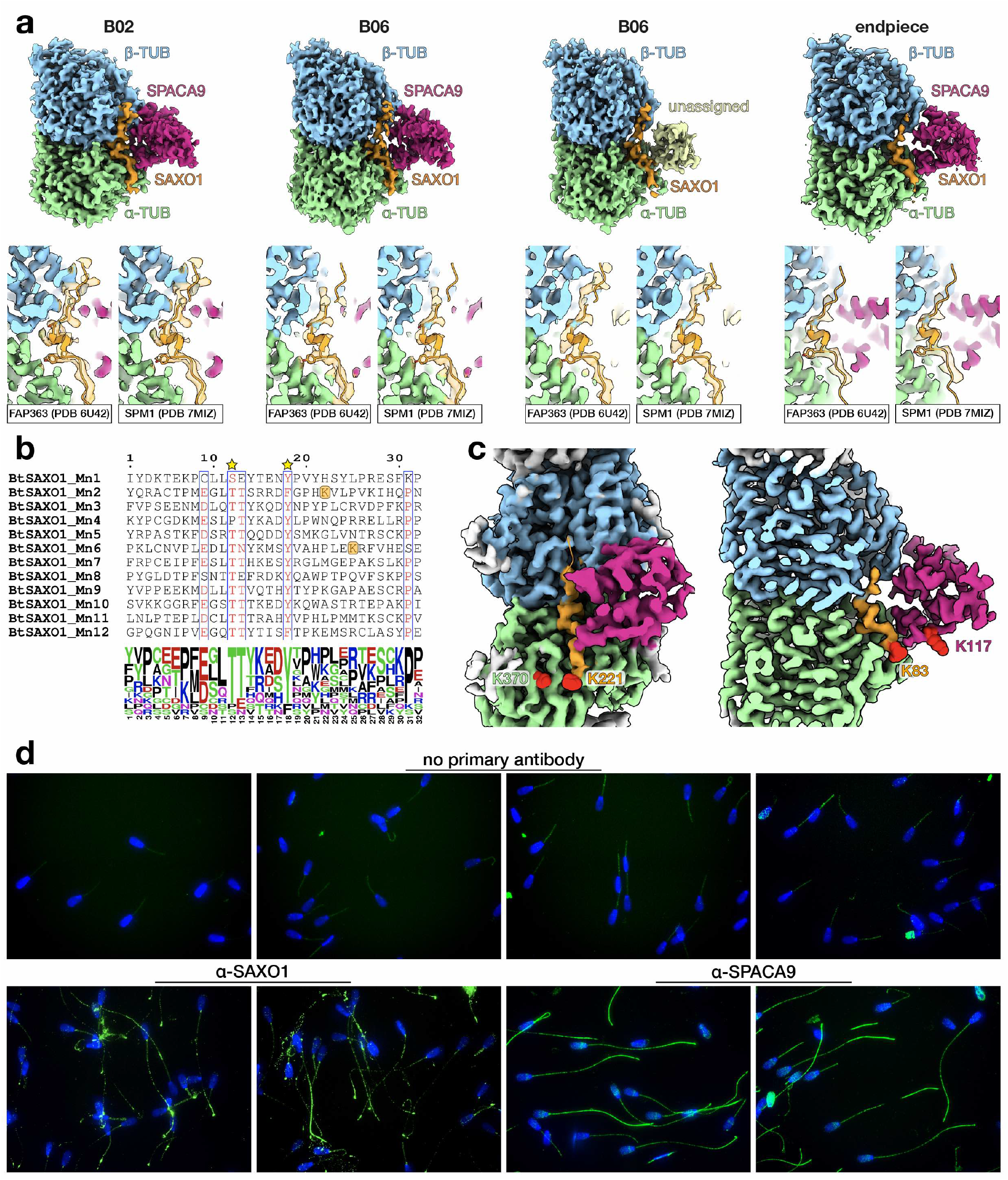
Additional data supporting the assignment of SAXO1 and SPACA9. **(a)** Top panels: extracted density for individual tubulin dimers and associated MIPs from protofilaments B02 and B06, and from the endpiece. Bottom panels: atomic models of the Mn motifs of FAP363 from *Chlamydomonas* DMTs ^5^ and SPM1 from *Toxoplasma* cortical MTs ^30^ fitted into the density for the filamentous MIP in bovine sperm B-tubules and endpiece DMTs. **(b)** Sequence alignment of the 12 Mn motifs within *Bos taurus* SAXO1 shows the consensus Thr and Tyr residues (stars) demarcating the short helix that binds at the alpha/beta-tubulin interface. Orange squares mark lysine residues (K83 and K221) that cross-linked to SPACA9 and alpha-tubulin respectively. **(c)** In-cell cross-linking mass spectrometry detects interactions between SAXO1 and alpha tubulin (left); and between SAXO1 and SPACA9 (right). Red spheres indicate the cross-linked lysine residues. **(d)** Immunofluorescence confirms that both SAXO1 and SPACA9 are present in bovine sperm flagella.

The mammalian homologs of FAP363 and SPM1 are the stabilizer of axonemal microtubules proteins, SAXO1 and SAXO2 ^32^. Our proteomics data show that SAXO1 is abundant in bovine sperm while no unique peptides for SAXO2 were detected **(Table S2)**. Furthermore, our in-cell cross-linking mass spectrometry (XL-MS) data show cross-links between a lysine close to an Mn motif of SAXO1 (Lys221) and Lys370 in the S9-S10 loop of α-tubulin, which is near the SAXO1 binding site **(Fig. 10b-c, Table S3)**. We also confirmed that SAXO1 is present in the bovine sperm flagellum using immunofluorescence **(Fig. 10d)**, which is consistent with its known presence in human sperm flagella based on immunofluorescence and immunogold labelling ^32^. Together, these lines of evidence suggest that the filamentous MIP in the B-tubule is SAXO1. Unsharpened maps suggest that SAXO1 binds longitudinally along individual protofilaments **(Fig. 8c, arrowheads)**, similar to SPM1. However, we note that SAXO1 has 12 Mn repeats, which means that it could have a 96-nm periodicity when fully extended. The register of SAXO1 Mn motifs could not be determined from the density alone, nor could the small N-or C-terminal domains be resolved. More detailed analysis of the SAXO1 binding mode – and the binding modes of different Mn motif proteins – is an important target for future work.

The second MIP (magenta) – consists of a bundle of alpha helices that binds across the intradimer α/β-tubulin interface as well as between adjacent protofilaments **(Fig. 8b,e**). Based on side chain densities, we identify this MIP as sperm acrosome-associated 9 (SPACA9). Further supporting this identification, the high-confidence AlphaFold structure of SPACA9 is a bundle of alpha-helices whose topology precisely matches the density profile of the MIP, barring a C-terminus that is predicted to be disordered and for which AlphaFold confidence scores are lower **(Fig. S6)**. Every 8-nm, there are four copies of SPACA9 that each bind to the inter-protofilament regions of B02-B06 **(Fig. 8a)**. Specifically, SPACA9 interacts with both α- and β-tubulin within a dimer while also interacting with the M-loop of α-tubulin from the neighboring protofilament **(Fig. 8e)**. Every 48-nm, one additional copy of SPACA9 binds between tubulin protofilaments B06/B07 **(Fig. 8a)**. The B06/B07 interface is normally occluded by CFAP45, but once every 48-nm CFAP45 curves away from the interface, freeing a binding site for SPACA9 **(Fig. 8d, arrowheads)**.

Every 48-nm there is another MIP (pale green) that binds close to the B06/B07 SPACA9 binding site **(Fig. 8a)**. Unlike SPACA9, this as-yet-unidentified MIP does not seem to bind tubulin in protofilament B07, and instead seems to recognize CFAP45 because at its binding site CFAP45 still occludes the B06/B07 inter-protofilament interface **(Fig. 8d)**. Nevertheless, this arrangement explains the varying lengths of the “ladder rungs” observed in subtomogram averages from mammalian sperm **(Fig. 8a, 9a panels i to iii)**, i.e. every 48-nm repeat has one long rung (covering B02-B07) followed by three short rungs (covering B02-B06), followed again by one long rung and finally by one short rung. As discussed above, the two long rungs are not identical – one is formed by the unidentified MIP **(Fig. 8a, 9a panel i)** and the other by an additional copy of SPACA9 **(Fig. 8a, 9a panel iii)**. Intriguingly, subtomogram averages from mouse sperm appear to have fewer MIPs in this region **(Fig. 9c)**, having shorter shelf-like protrusions rather than the extensive ladder-like rungs seen in pig and horse ^12,33^.

### Endpiece singlet microtubules share MIPs with the B-tubule of the axonemal doublets

At the end of the principal piece, the characteristic structure of the axoneme is lost and DMTs transition into singlet MTs (SMTs) ^12,34^. The region comprising mainly SMTs ensheathed by the plasma membrane is known as the endpiece. Cryo-ET and subtomogram averaging resolved discontinuous intraluminal spirals in endpiece SMTs from the sperm of several mammals ^12,19^. Because these discontinuous spirals resemble the ladder-like ridges in the B-tubule, we sought to determine whether the two structures are made up of similar MIPs. To address this, we determined high-resolution structures of endpiece SMTs from whole bovine sperm **(Fig. 11)**. To best preserve the native structure of endpiece SMTs, we imaged whole untreated bovine sperm and searched for endpieces that had spontaneously frayed during sample preparation, likely due to shear forces on the membrane either from centrifugation, pipetting, or blotting **(Fig. 11a)**.

**Fig. 11.**
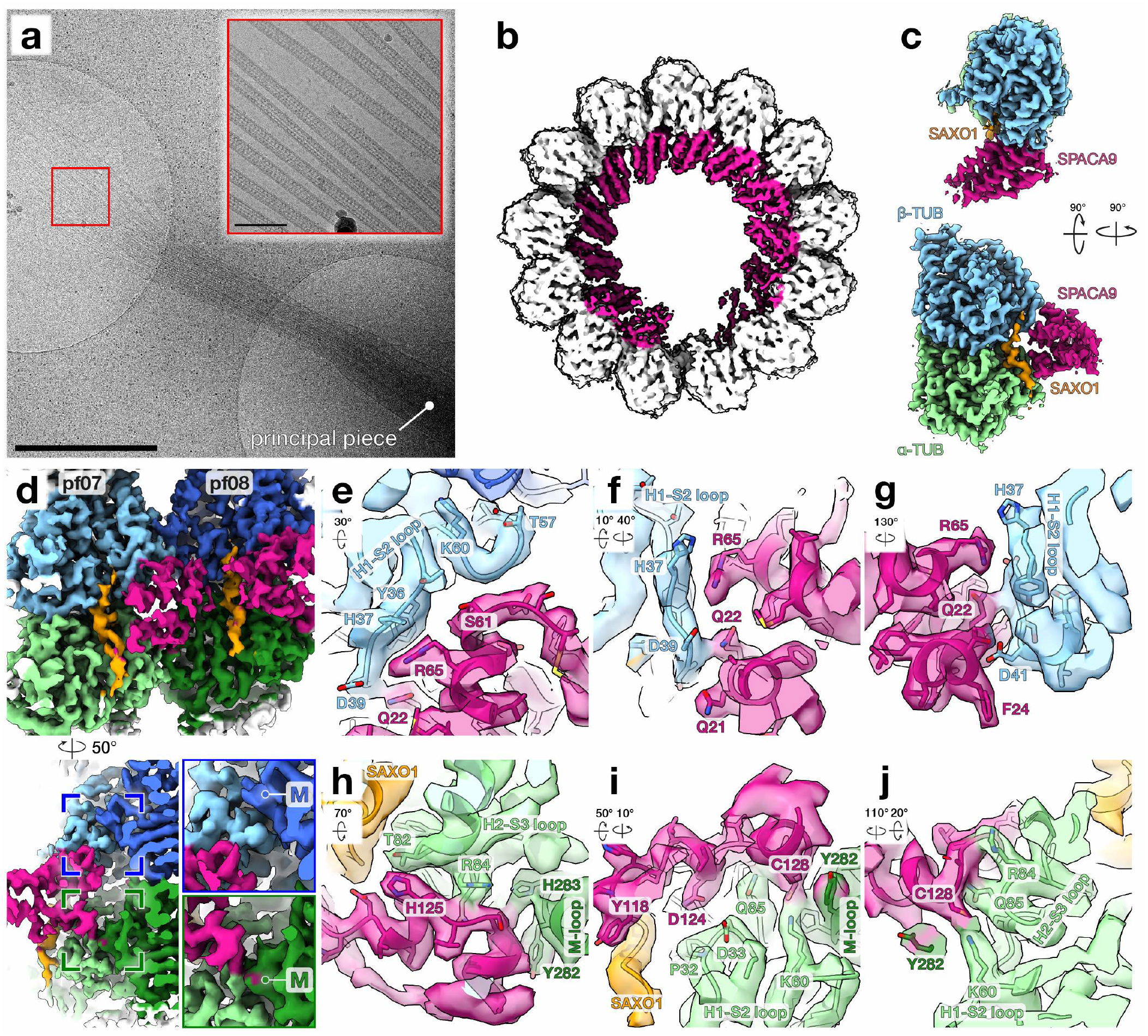
Cryo-electron microscopy of singlet microtubules in frayed endpieces of whole bovine sperm. **(a)** Cryo-EM image of a frayed sperm endpiece on a holey carbon grid. Note how the fibrous sheath, which demarcates the principal piece, remains intact (along with, in this case, the plasma membrane around it). Scale bar: 1 μm. Inset: High-magnification cryo-EM image of endpiece singlet microtubules (SMTs) captures a ladder-like luminal structure with an ∼8-nm periodicity. Scale bar: 100 nm. **(b)** Cryo-EM reconstruction of endpiece SMTs (lowpass-filtered to 5 Å) reveals that the discontinuous intraluminal spirals consist of predominantly alpha-helical microtubule inner proteins (MIPs) projecting into the lumen. **(c)** Map of the asymmetric unit obtained after symmetry expansion reveals that SPACA9 and SAXO1 are also MIPs in endpiece SMTs. **(d)** Top panel: Luminal view of the inter-protofilament interface obtained from local refinement of protofilaments 7 and 8. Bottom panel: Rotated and clipped view showing how SPACA9 interacts with both α and β-tubulin, along with the M-loop of α-tubulin from the neighboring protofilament (green box) but not the M-loop of the neighboring β-tubulin (blue box). **(e-g)** Close-up views of interactions between SPACA9 and the H1-S2 loop of β-tubulin. **(h-j)** Close-up views of interactions between SPACA9 and the H1-S2 and H2-S3 loops of α-tubulin. Note how SPACA9 also interacts with SAXO1 (i) and the M-loop of α-tubulin from the neighboring protofilament (h,i). Arrows under panel labels in (e-j) indicate how the zoomed-in views are rotated relative to the top panel in (d).

We resolved C1 structures of bovine sperm microtubules at global nominal resolutions of ∼4.3 Å **(Fig. 11b, Fig. S7b)**. Mammalian sperm endpiece SMTs are 13-protofilament microtubules with a rise of 9.57 Å and a twist of -27.71° as estimated by relion_helix_toolbox. To improve the cryo-EM density towards identifying MIPs in endpiece SMTs, we performed symmetry expansion, protofilament-based subtraction, 3D classification, and local refinement, similar to the strategy described in ^35^. This resulted in a ∼3.5-Å map with well-resolved side chain densities, which greatly facilitated interpretation, protein identification, and model building **(Fig. S7b-e)**. Our cryo-EM maps allow us to identify the MIPs as SPACA9 and SAXO1 **(Fig. 11c, 10-S7)**, which are shared between endpiece SMTs and the B-tubule of DMTs. To analyze how SPACA9 interacts with tubulin, we fit models of the asymmetric unit derived from the symmetry-expanded maps into locally-refined maps of protofilaments 07 and 08 **(Fig. 11d-g)**. Within a dimer, SPACA9 makes contacts with the H1-S2 loop of β-tubulin **(Fig. 11e-g,**) and interacts with both the H1-S2 and H2-S3 loops of α-tubulin **(Fig. 11h-j)**.

The SPACA9 binding mode in endpiece SMTs is similar to that of SPACA9 in the B-tubule of axonemal DMTs; specifically, SPACA9 interacts with the M-loop of α-tubulin in the neighboring protofilament **(Fig. 11h-i)** but is too far to interact with the corresponding M-loop of the neighboring β-tubulin **(Fig. 11d, bottom panel)**. SPACA9 also interacts close to the Mn motif of SAXO1 **(Fig. 11i)**, which is consistent with cross-links identified between SPACA9 Lys117 and SAXO1 Lys83, which is close to an Mn motif **(Fig. 10b,c**). The difference in protofilament curvature between endpiece SMTs and the B-tubule means that the rotation between neighboring SPACA9 molecules also differs between SMTs and DMTs **(Fig. 12a)**, reflecting the hinge-like flexibility of the M-loops at the inter-protofilament interface **(Fig. 12b)**.

**Fig. 12.**
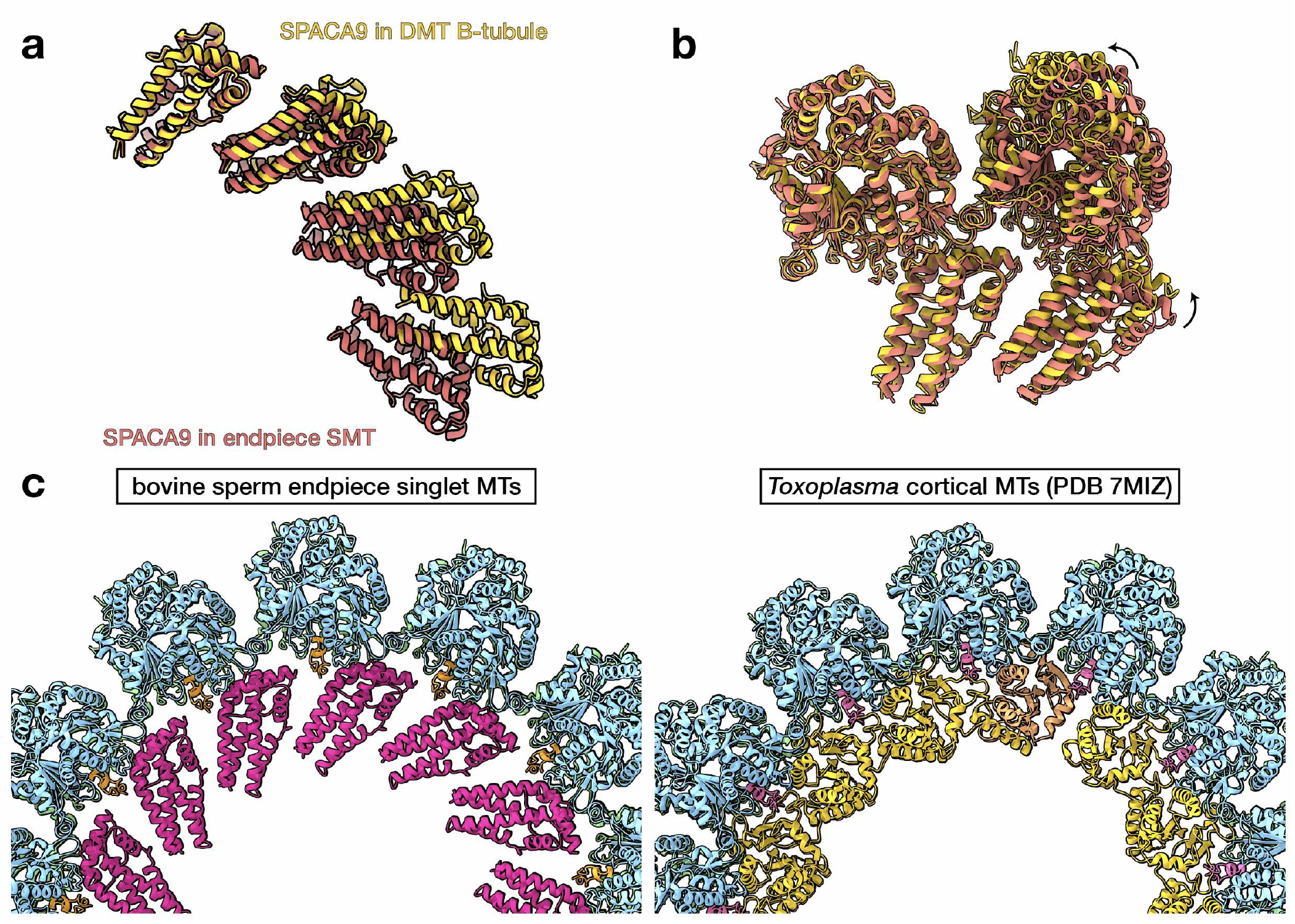
Comparing sperm endpiece singlet microtubules with sperm doublet microtubules and *Toxoplasma* cortical microtubules. **(a)** SPACA9 is shared between endpiece singlets and the B-tubule of axonemal doublets, and the SPACA9 arrays reflect underlying differences in protofilament curvature between the two microtubules. **(b)** Overlay of two neighboring α/β-tubulin/SAXO1/SPACA9 units from the DMT B-tubule (gold) and from the endpiece SMT (coral). Note the hinge-like motion centered on the M-loops. **(c)** Atomic models of bovine sperm endpiece microtubules and *Toxoplasma gondii* cortical microtubules (PDB 7MIZ) 30 show similar MIP arrangements. SAXO1 in mammalian sperm shares a binding mode with SPM1 in *Toxoplasma*, while SPACA9 in mammals partly shares a binding interface with Trxl1/2 in *Toxoplasma*. While SAXO1 and SPM1 are members of the same protein family, SPACA9 and Trxl1/2 are structurally unrelated.

## Discussion

In this study, we used cryo-EM to solve structures of axonemal DMTs and endpiece SMTs from mammalian sperm, revealing extensive ornamentation by sperm-MIPs. Ciliary MIPs have been shown to stabilize microtubules ^36,37^, so the elaborate interaction networks formed by sperm-MIPs likely serve to further reinforce the microtubule lattice itself against the large mechanical stresses involved in bending the long, stiff flagella of mammalian sperm. In particular, the outer dense fibers make the axoneme more rigid while also increasing its effective diameter and thus concomitantly increasing bending torque, entraining more dyneins with every bend ^15^. Because interactions between protofilaments are weaker than interactions between tubulin monomers along a protofilament ^38,39^, protofilaments can slide relative to one another, conferring shearability onto the microtubule. Protofilament-bridging MIP networks could act as structural braces to decrease inter-protofilament shearing, thus increasing microtubule bending stiffness.

Whereas axoneme structure has been studied extensively, the distal region of the cilium – called the endpiece in sperm flagella – is much more poorly understood ^40^. The endpiece consists only of SMTs and lacks dynein motors and other axonemal complexes. Mathematical modelling suggests that the presence of an inactive non-bending distal region has important effects on the ciliary waveform ^41^. Our structures show conclusively that endpiece SMTs share MIPs with the B-tubule of axonemal DMTs, specifically the MIPs SAXO1 and SPACA9. Given that endpiece SMTs are derived from both the A- and B-tubules of axonemal DMTs in mammalian sperm ^12,34^, all the MIPs in the A-tubule must at some point be replaced with SPACA9 and SAXO1. Precisely how this transition occurs and what signals it are important questions for future work.

From a structural perspective, endpiece SMTs in mammalian sperm are very similar to cortical MTs from the parasite *Toxoplasma gondii* **(Fig. 12c)**. Both microtubules are stabilized by filamentous Mn motif-containing MIPs that bind longitudinally along protofilaments (SPM1 in *Toxoplasma* and SAXO1 in sperm). Both microtubules are further stabilized by MIPs that bind across protofilaments, although in this case the MIPs (TrxL1/2 in *Toxoplasma* and SPACA9 in sperm) are structurally-unrelated. This points to similar architectural principles for stabilizing microtubules from the lumen, which are likely to be more widespread given that similar protofilament-spanning MIP densities have also been observed in *Plasmodium* gliding forms ^42^.

Beyond providing insights into sperm biology, our cryo-EM structures, together with recent maps of DMTs from bovine respiratory cilia, paint a concrete and detailed portrait of the molecular and architectural diversity of motile cilia across different cell types within an organism. However, while the Human Protein Atlas reports expression of many sperm-MIPs as testis-enriched ^24^, a number have been reported in other ciliated cell types as well. For example, PPP1R32 was detected in ependymal cilia in brain ventricles ^43^, while CFAP77 and SPACA9 were detected in ciliated airway cells ^44^. It is possible that these MIPs are not strictly sperm-specific, but that their expression along or around the sperm axoneme is more widespread than in other ciliated cell types. Indeed, PPP1R32, CFAP77, SPACA9, and TTBP were detected in proteomics of the same bovine tracheal axoneme samples used to obtain cryo-EM maps in ^6^. It is possible that these MIPs were not resolved in cryo-EM maps of respiratory cilia because their expression is restricted to only a few of the nine DMTs or to a shorter region along the axoneme. Consistent with this hypothesis, subtomogram averages obtained from different regions of bovine respiratory cilia showed ladder-like ridges resembling those formed by SPACA9, but only in the transition zone ^14^.

Our study also raises fascinating evolutionary questions – how does ornamentation with extra sperm-MIPs correlate with the appearance of flagellar accessory structures? Because accessory structures are thought to prevent buckling instabilities when sperm swim through viscous media ^18^, how do sperm-MIPs likewise vary across internally- and externally-fertilizing species? Sperm from external fertilizers such as sea urchin ^9^ and zebrafish ^45^ do not appear to have as many MIPs as their mammalian counterparts ^12,20,21^. The natural diversity of sperm form ^46^ thus presents a unique opportunity to understand the evolution and diversification of a core cellular machine; with the approaches we describe in this work, this question can now be addressed from a truly molecular level, integrating perspectives from genomic, proteomic, and structural methods.

For example, our structures raise questions about the nature and evolution of Tektin-5. While all other Tektins form filaments through similar intermolecular interactions, Tektin-5 adopts a range of conformations in the sperm DMT. Tektin-5 appears to be testis-specific, and recent bioinformatics analyses ^47^ concluded that Tektin-3 and Tektin-5 arose from duplication of the Tektin-3/5 gene in early vertebrates; interestingly, Tektin-5 was subsequently lost in several species of teleost fish, including the zebrafish *Danio rerio*, and amphibians, including the frog *Xenopus laevis*. This may explain why the A-tubule in zebrafish sperm DMTs appear more “empty” than their mammalian counterparts ^45^. Loss of Tektin-5 does not appear to correlate with fertilization mode, as it was lost in both the externally-fertilizing *X. laevis* and the internally-fertilizing *Notophthalmus viridiscens* (Eastern newt), but retained in the externally-fertilizing fish species *Lepisosteus oculatus* (spotted gar) and *Clupea harengus* (herring) ^47^. These observations raise questions about the functional roles of Tektin-5. Mice deficient in Tektin-2 or -4 are infertile or subfertile respectively ^48,49^, and mice lacking Tektin-3 have reduced sperm motility and forward progression despite showing normal fertility ^50^. While the effects of Tektin-1 deficiency on sperm have not been explicitly tested, loss of Tektin-1 causes ciliary defects in zebrafish ^51^. These phenotypes and our structures would predict that disrupting Tektin-5 would affect sperm motility and male fertility; however, there is currently no literature on the effects of genetically-ablating Tektin-5, which is clearly an important target for further research into sperm motility.

In contrast, mouse knockouts for several MIPs identified in our cryo-EM structures have in fact been reported in the literature. These include TEX43 ^52^, C7ORF31 ^53^, SPACA9 ^54^, SAXO1 ^55^, SMRP1, and TEPP ^56^. In all cases, knockout mice were fertile; this is not necessarily surprising given that a certain degree of robustness can be expected for essential processes such as those related to fertilization and reproduction. Such robustness has previously been observed in motile cilia, where *Chlamydomonas* mutants for three inner junction proteins FAP20, FAP45, and FAP52 nonetheless retained some attached inner junctions ^36^. Furthermore, the absence of a male infertility phenotype does not exclude the possibility of more subtle effects on motility; for instance, sperm from TEX43-deficient mice did show reduced velocity parameters ^52^. It is also possible that infertility only results when several sperm-MIPs are knocked out simultaneously. By identifying many of the sperm-MIPs, our structures now serve as valuable resources for targeted functional and genetic studies aimed at dissecting the roles of individual proteins on sperm motility. In a similar vein, our work could also provide a structural framework for understanding male infertility, which is on the rise globally ^57,58^ yet remains unexplained in many cases ^59^.

## Acknowledgements

The authors thank Dr. H Henning and A Rijneveld of the Utrecht University Veterinary Faculty for providing bovine sperm. The authors acknowledge Ingr. CTWM Schneijdenberg and JD Meeldijk for managing and maintaining the Utrecht University EM Facility. The authors thank Dr. M Vanevic for computational support, and Dr. SC Howes for valuable discussions on data collection and processing strategies. The authors also thank Prof. A Akhmanova for critical reading of the manuscript. This project benefitted from access to the Netherlands Centre for Electron Microscopy (NeCEN) with support from operator Dr. W Noteboorn. RZC, JFH, and AJRH acknowledge support from the NWO X-omics Road Map program project 184.034.019. This work was funded by NWO Start-Up Grant 740.018.007 to TZ.

## Author Contributions

MRL and MCR prepared sperm samples. MRL and MCR collected and processed cryo-EM data. MRL and TZ analyzed data and built atomic models. RZC and JH performed proteomics and cross-linking mass spectrometry experiments and corresponding data analysis, supervised by AJRH. MRL and TZ wrote the manuscript, and all authors contributed to revisions.

## Materials and Methods

### Cryo-EM of Doublet Microtubules

#### Sample preparation

Frozen bovine semen straws from the Utrecht University Veterinary Faculty were thawed rapidly in a 37°C water bath. Sperm cells were washed twice in 1X Dulbecco’s phosphate-buffer saline (DPBS, Sigma) and counted. To remove membranes and the mitochondrial sheath ^22^, sperm were diluted to ∼1-2 × 10^5^ cells/mL in demembranation buffer (20 mM Tris-HCl pH 7.9, 132 mM sucrose, 24 mM potassium glutamate, 1 mM MgSO_4_, 1 mM DTT, and 0.1% Triton X-100), frozen at -20°C for 48-96 h, then thawed. To induce sliding disintegration, ATP (Sigma) was added to a final concentration of 1 mM and the solution incubated for 10-15 min at room temperature. Axoneme disintegration was verified under an inverted light microscope.

Approximately 4 μL of disintegrated sperm suspension was applied to glow-discharged Quantifoil R 2/1 200-mesh holey carbon grids, which were then blotted from the back for ∼5-6 s using a manual plunger (MPI Martinsried, Germany). Grids were plunged into a liquid ethane/propane mix (37% ethane) cooled to liquid nitrogen temperatures. Grids were clipped into Autogrids (ThermoFisher) and stored under liquid nitrogen until imaging.

#### Cryo-electron microscopy

A total of five datasets were collected for this study, from a total of 7 grids from 3 sperm straws. The first four were collected on a Talos Arctica (ThermoFisher) operated at 200 kV and equipped with a GIF Quantum K2. The last was collected on a Titan Krios (ThermoFisher) operated at 300 kV and equipped with a BioQuantum K3. For all datasets, the energy filter was operated in zero-loss mode with a 20-eV slit width. All images were collected in super-resolution mode with a total electron fluence of ∼48-50 e-/Å^2^, with ∼1-1.1 e-/Å^2^ per frame. Acquisition areas were identified manually and images were collected semi-automatically in SerialEM ^60^. Frames were motion-corrected on-the-fly using Warp ^61^ to monitor data quality during the session.

#### Cryo-EM data processing

All data processing was performed in Relion 3.1 ^62^ based on strategies described in ^5,6^. Specific details of processing are reported in **Table S1**. The processing workflow is summarized in **Fig. S1**. Super-resolution frames were binned twice, motion-corrected, and dose-weighted using Relion’s implementation of MotionCor2. The contrast transfer function (CTF) was estimated using CtfFind. Microtubules were picked manually and particles were extracted every ∼82 Å (the length of a tubulin dimer). For initial alignments, twice-binned particles were extracted with a box size of 336 (∼700 Å, encompassing ∼9 tubulin dimers).

Global alignment parameters were first determined for the 8-nm particles through a C1 helical auto-refinement in Relion 3.1. The cryo-EM map of the doublet microtubule (DMT) from bovine respiratory cilia (EMD-24664) ^6^ filtered to 30-Å was used as an initial reference. To enrich for fully-decorated DMTs, tubulin signal was subtracted and 3D classification performed with a mask covering MIPs bound to protofilaments B02-B06. Classes with well-resolved density were selected for further processing. Particles offset by 4 nm were also identified at this stage and shifted back into register with the majority class. After checking for duplicate particles, the remaining particles were entered to a 3D auto-refinement, yielding a map of the 8-nm repeat.

To retrieve the 16- and 48-nm repeats, tubulin signal was subtracted and 3D classification performed first with a mask covering MIPs at the inner junction, then with a mask covering MIPs at the seam of the A-tubule. The resulting 48-nm particles were re-extracted with a box size of 672 and refined. After CTF refinement (magnification anisotropy correction, followed by per-particle defocus estimation and aberration correction) and Bayesian polishing, the nominal overall resolution of the final map was ∼4 Å. To further improve resolution, we performed local refinements, each with a cylindrical mask covering 2-3 protofilaments and extending along ∼1/3 of the 48-nm repeat **(Fig. S1)**. We individually post-processed and sharpened the 30 locally-refined maps, then generated a composite map using the *fit in map* and *volume maximum* commands in ChimeraX ^63^. We used the same strategy to generate composite half-maps to assess overall resolution, estimated at ∼3.7 Å, and local resolutions, estimated to reach ∼3 Å in the core of the A-tubule.

#### Cryo-EM data processing

Map interpretation was guided by the atomic model of the DMT from bovine respiratory cilia (PDB 7RRO) ^6^. The positions of tubulin dimers and MIPs were manually adjusted by rigid body fitting in ChimeraX, followed by real-space refinement in Coot and in Phenix. The α-tubulin sequences were mutated to match the most abundant isoform identified in sperm (TUBA3); β-tubulin sequences already corresponded to the most abundant sperm isoform (TUBB4B). Tubulin C-terminal tails and the α-K40 loop were manually built based on the density map. Models for MIPs already present in respiratory cilia were likewise truncated or extended based on the density.

Unknown MIPs were identified using a workflow based on the findMySequence program ^23^. Briefly, starting poly-Ala models were manually built into the map using ‘Place Helix Here’ or ‘Place Strand Here’ tools in Coot, manually extended when the density permitted, then real-space refined in Coot. The findMySequence program was then used to estimate side chain probabilities and to query a database consisting of the 1500 most abundant proteins in bovine sperm (see section “Mass spectrometry”). Once confident protein identities were obtained, findMySequence was also used to assign sequences to the query poly-Ala model. The models were then manually extended using the positions of bulky side chains as guideposts.

This workflow was first validated on known MIPs, like RIBC2, and could reliably distinguish between closelyrelated paralogs like Tektins1-4 **(Fig. S3a)**. Backbone traces were generally sufficient for findMySequence to identify MIPs of varying lengths and folds **(Fig. S3b)**; however, in the case of DUSP21, running findMySequence directly on traceable secondary structure elements did not yield a hit. Querying the DALI server ^64^ returned dual-specificity phosphatase domains as hits. The top hit (PDB 5Y16) ^65^ was mutated to polyAla and fitted into the density map of the unknown MIP. Running findMySequence on this model returned DUSP21 as a confident hit **(Fig. S3c)**. For SPACA9, findMySequence could assign protein identity, but it was initially difficult to fully trace the backbone **(Fig. S3d)**. The high-confidence AlphaFold2 prediction ^66^ for SPACA9 was thus used to facilitate model building.

For the first round of real-space refinement in Phenix, the 48-nm repeat was divided into several PDB files corresponding to each MIP or to each protofilament of the doublet. Each model was refined individually, then manually inspected in Coot. For subsequent rounds, PDB files were merged into a single model of the 48-nm repeat, which was refined as a whole against the composite map. The refined structures were used to measure inter-protofilament angles and inter-dimer distances, which were estimated in ChimeraX using the align and distance commands respectively.

### Cryo-EM of Endpiece Singlet Microtubules

#### Sample preparation

Frozen bovine semen straws from the Utrecht University Veterinary Faculty were thawed rapidly in a 37°C water bath. Sperm cells were washed twice in 1X DPBS (Sigma) and counted. Sperm concentration was adjusted to ∼1-3 × 10^6^ cells/mL. Approximately 4 μL of untreated sperm suspension was applied to glow-discharged Quantifoil R 2/1 200-mesh holey carbon grids, which were then blotted from the back for ∼4-6 s using a manual plunger (MPI Martinsried, Germany). Grids were plunged into either liquid ethane or a liquid ethane/propane mix (37% ethane) cooled to liquid nitrogen temperatures. Grids were clipped into Autogrids (ThermoFisher) and stored under liquid nitrogen until imaging.

#### Cryo-electron microscopy

Cryo-EM data for endpiece singlet microtubules was collected over 7 imaging sessions from a total of 8 grids from 3 sperm straws. Grids were imaged on a Talos Arctica (ThermoFisher) operated at 200 kV. The microscope was additionally equipped with a GIF energy filter (Gatan), which was operated in zero-loss mode with a 20-eV slit width. All images were collected in counting mode on a K2 Summit direct electron detector (Gatan) with a total electron fluence of ∼48-50 e-/Å^2^, with ∼1-1.1 e-/Å^2^ per frame. Acquisition areas were identified manually and images were collected semi-automatically in SerialEM. Frames were motion-corrected on-the-fly using Warp to monitor data quality during the session.

#### Cryo-EM data processing

All data processing was performed in Relion 3.1 unless otherwise noted. Specific details of processing are reported in **Table S1**. The processing workflow is summarized in **Fig. S7**. Frames were motioncorrected and dose-weighted using Relion’s implementation of MotionCor2. The contrast transfer function (CTF) was estimated using gctf. Microtubules were picked manually and particles were extracted every ∼82 Å (the length of a tubulin dimer) with a box size of 432 (∼587 Å, encompassing ∼7 dimers).

An initial C1 helical auto-refinement was performed using a ∼20-Å subtomogram average of pig endpiece singlet microtubules (EMD-12068) ^12^ as a reference, resulting in a ∼4.8-Å map. To improve particle alignments towards an improved C1 reconstruction, the microtubule Relion-based pipeline (MiRPv2) was used to perform rotation angle smoothing, XY shift smoothing, and seam correction ^67,68^. Aligned, seam-corrected particles were then subjected to a round of C1 helical auto-refinement with restricted searches and a mask encompassing the central 40% of the microtubule. This resulted in a C1 reconstruction at ∼4.6-Å resolution, which was improved to ∼4.3-Å after two rounds each of CTF refinement and Bayesian polishing. Local refinements of protofilaments 7/8 improved resolution to ∼4-Å.

To improve MIP densities, symmetry-expansion was performed on the dataset based on helical parameters estimated from relion_helix_toolbox (rise = 9.57 Å, twist = -27.7°) ^35^. Particle subtraction was first run with a mask covering four tubulin dimers of the “good” protofilament opposite the seam (protofilament 7), followed by 3D classification of the resulting particles without image alignment. The class with the best MIP density was selected and entered into a local refinement with a mask on two central tubulin dimers. This resulted in a ∼3.5-Å map with well-resolved side chain densities that we used for protein identification and model building.

#### Model building and refinement

Modelling was based on the cryo-EM map of one asymmetric unit (one copy each of alpha- and beta-tubulin, SPACA9, and an Mn motif of SAXO1) obtained after symmetry expansion. To model the tubulin dimer, a model of porcine tubulin from PDB 3JAS ^69^ was used as a starting model and mutated to match the appropriate bovine tubulin sequences. Tubulin isoforms (TUBA3 and TUBB4B) were chosen based on abundance from proteomics data and on side chain density at distinguishing residues. To model the Mn motif of SAXO1, SPM1 from PDB 7MIZ ^30^ was used as a starting model. Residues were then mutated to match the (arbitrarily chosen) sixth Mn motif of bovine SAXO1. To model SPACA9, an AlphaFold model was initially used, and residues not resolved in the map were deleted from the C-terminus. The combined model was real-space refined in Phenix, manually adjusted in Coot, and finally real-space refined in Phenix.

### Mass spectrometry

#### Cross-linking, lysis, digestion, and peptide fractionation

All proteomics and cross-linking mass spectrometry experiments were performed essentially according to ^70^ on bovine sperm prepared as described above. The sperm cells were resuspended in 540 μL of PBS and supplemented with DSSO (ThermoFisher Scientific) to a final concentration of 1 mM. The reaction was incubated for 30 min at 25°C with 700 rpm shaking in a ThemoMixer C (Eppendorf) and subsequently quenched for 20 min by adding Tris-HCl (final concentration 50 mM). Cells were centrifuged at 13800×g for 10 min at 4°C, and the supernatant was replaced with lysis buffer. Cells were resuspended in 1 mL of lysis buffer (100 mM Tris-HCl pH 8.5, 7 M Urea, 1% Triton X-100, 5 mM TCEP, 30 mM CAA, 10 U/ml DNase I, 1 mM MgCl_2_, 1% benzonase (Merck Millipore, Darmstadt, Germany), 1 mM sodium orthovanadate, phosphoSTOP phosphatases inhibitors, and cOmpleteTM Mini EDTA-free protease inhibitors) and lysed with the help of sonication (2 minutes with UP100H from Hielscher at 80% amplitude). The proteins were then precipitated and resuspended in digestion buffer (100 mM Tris pH 8.5, 1% sodium deoxycholate [Sigma-Aldrich], 5 mM TCEP, and 30 mM CAA). Trypsin and Lys-C proteases were added to a 1:25 and 1:100 ratio (weight/weight), respectively, and protein digestion performed overnight at 37°C shaking at 1300rpm on ThemoMixer C. Peptides were then desalted with Oasis HLB plates (Waters) and fractionated with an Agilent 1200 HPLC pump system (Agilent) coupled to a strong cation exchange (SCX) separation column (Luna SCX 5 μm to 100 Å particles, 50×2 mm, Phenomenex), resulting in 24 fractions. Each fraction was then desalted with OASIS HLB plate.

#### Liquid chromatography with mass spectrometry

Before injecting each SCX fraction, 1,000 ng of peptides from each biological replicate were first injected onto an using an Ultimate3000 high-performance liquid chromatography system (ThermoFisher Scientific) coupled online to an Orbitrap HF-X (ThermoFisher Scientific). For this classical bottom-up analysis, we used the following parameters as in ^71^: Buffer A consisted of water acidified with 0.1% formic acid, while buffer B was 80% acetonitrile and 20% water with 0.1% formic acid. The peptides were first trapped for 1 min at 30 μl/min with 100% buffer A on a trap (0.3 mm by 5 mm with PepMap C18, 5 μm, 100 Å; ThermoFisher Scientific); after trapping, the peptides were separated by a 50-cm analytical column packed with C18 beads (Poroshell 120 EC-C18, 2.7 μm; Agilent Technologies). The gradient was 9 to 45% B in 95 min at 400 nl/min. Buffer B was then raised to 55% in 10 min and increased to 99% for the cleaning step. Peptides were ionized using a spray voltage of 2 kV and a capillary heated at 275°C. The mass spectrometer was set to acquire full-scan MS spectra (350 to 1400 mass/charge ratio) for a maximum injection time of 120 ms at a mass resolution of 120,000 and an automated gain control (AGC) target value of 3 × 10^6^. Up to 25 of the most intense precursor ions were selected for MS/MS. HCD fragmentation was performed in the HCD cell, with the readout in the Orbitrap mass analyser at a resolution of 15,000 (isolation window of 1.4 Th) and an AGC target value of 1 × 10^5^ with a maximum injection time of 25 ms and a normalized collision energy of 27%.

The SCX fractions were analysed with same Ultimate HPLC and the same nano-column coupled on-line to an Orbitrap Lumos mass spectrometer (ThermoFisher Scientific). For these runs, we used same gradient and LC setting of bottom up data with specific MS settings for DSSO cross-links: survey MS1 Orbitrap scan at 120,000 resolution from 350 to 1,400, AGC target of 250% and maximum inject time of 50 ms. For the MS2 Orbitrap scan we used 30,000 resolution, AGC target of 200%, and maximum inject time of 118 ms for detection of DSSO signature peaks (difference in mass of 37.972 Da). The four ions with this specific difference were analysed with a MS3 Ion Trap scans at AGC target of 200%, maximum inject time of 200 ms for sequencing selected signature peaks (representing the individual peptides).

#### Data processing

Raw raw files obtained with classical bottom-up approach were analysed with MaxQuant version 1.6.17 with all the automatic settings adding Deamidation (N) as dynamic modification against the *Bos taurus* reference proteome (Uniprot version of 02/2021 with 37,512 entries). With this search, we were able to calculate intensity-based absolute quantification values and created a smaller FASTA file to use for analysis of cross-linking experiments. Raw files for cross-linked cells were analysed with Proteome Discoverer software version 2.5 (ThermoFisher Scientific) with the incorporated XlinkX node for analysis of cross-linked peptides as described by ^72^. Data were searched against the smaller FASTA created in house with “MS2_MS3 acquisition strategy”. For the XlinkX search, we selected full tryptic digestion with three maximum missed cleavages, 10 ppm error for MS1, 20ppm for MS2, and 0.5 Da for MS3 in Ion Trap. For modifications, we used static Carbamidomethyl (C) and dynamic Oxidation (M), Deamidation (N), and Met-loss (protein N-term). The crosslinked peptides were accepted with a minimum score of 40, minimum score difference of 4, and maximum false discovery rate set to 5%; further standard settings were used.

### Fluorescence microscopy

#### Immunofluorescence of SPACA9 and SAXO1 in bovine sperm flagella

Frozen bovine semen straws were thawed in a 37°C water bath. Thawed semen was diluted in DPBS and centrifuged at 500g for 10 min. The sperm pellet was washed twice with DPBS. The washed sperm suspension was diluted to a concentration of ∼5 × 10^6^ cells/mL. Approximately 150 μL of the suspension was pipetted into the wells of an 8-well ibidi μ-slide and left undisturbed for 30 min at RT to allow cells to settle. The sperm were then fixed with 4% formaldehyde in DPBS for 15 min at RT, then permeabilized with 0.1% Triton X-100 in DPBS for 15 min at RT. Cells were washed for 10 min in DPBS. Slides were then blocked with 1% BSA in DPBS for 1 h at RT. Sperm were then incubated with either anti-SAXO1 (HPA023899 from Atlas Antibodies, used at 2 μg/mL), anti-SPACA9 (HPA022243 from Atlas Antibodies, used at 6 μg/mL), or no primary antibody diluted in blocking buffer for 2 h at RT. After three 10-min washes with DPBS, sperm were incubated with 2° antibody (goat anti-rabbit IgG H+L AlexaFluor488 conjugate, 1/500) diluted in blocking buffer for 45 min at RT. Sperm were then counterstained with DAPI (1/1000 in DPBS) for 10 min at RT and finally washed thrice with DPBS. A few drops of Fluoroshield mounting medium were then applied to the wells and the slides stored at 4°C in the dark until imaging. Fluorescence microscopy was performed on a ThermoScientific CorrSight in spinning disk mode with a 63X 1.4-NA oil-immersion objective. Images were analyzed using Fiji v 2.0.0-rc-69/1.52p.

**Fig. S1.**
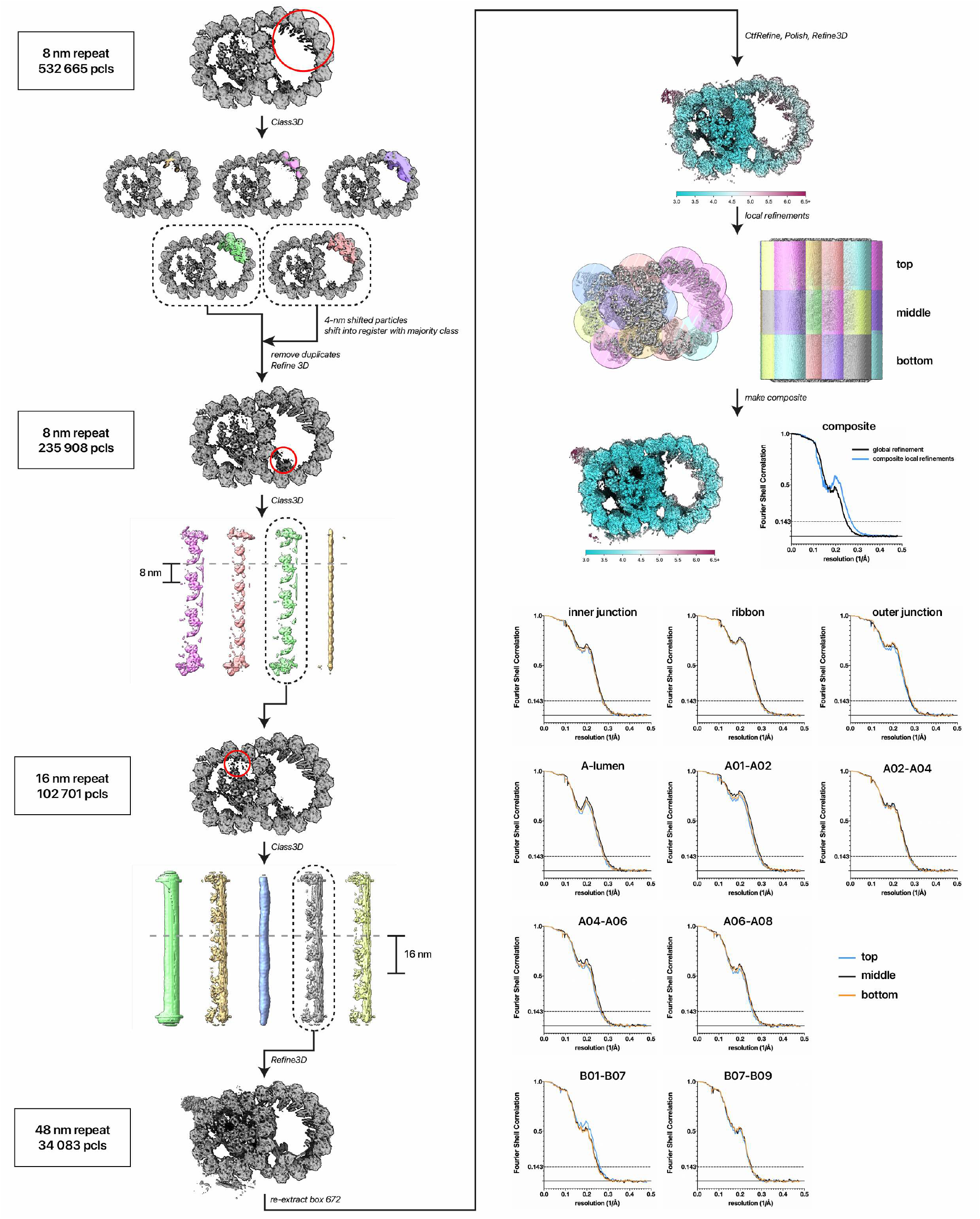
Processing workflow used to obtain a cryo-EM map of mammalian sperm DMTs. Microtubules were picked manually and particles were extracted every 8 nm. After classifying on B-tubule MIPs to enrich for good particles and fully-decorated DMTs, the 16- and 48-nm repeats were recovered by classifying on MIPs at the inner junction and the seam respectively. The 48-nm particles were re-extracted with a box size of 672, CTF-Refined, and polished. To improve resolution for model building, a series of 30 local refinements were performed using the masks shown in the figure. See Methods for more details.

**Fig. S2.**
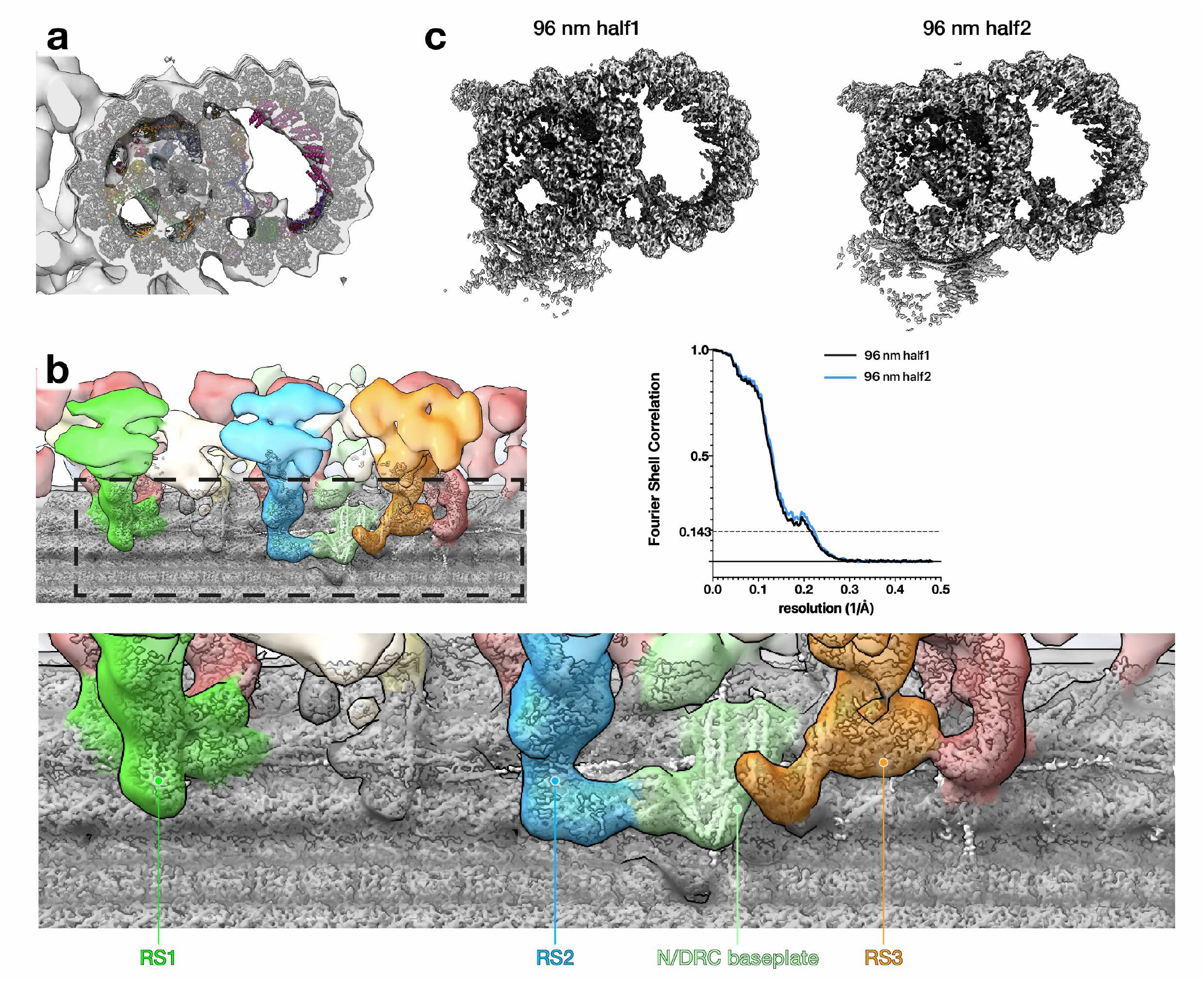
Reconstructing the 96-nm DMT repeat from mammalian sperm. **(a)** Fitting the atomic model of the 48-nm repeat into in situ subtomogram average of DMTs from pig sperm (EMD-12070) shows that all prominent densities are retained. **(b-c)** To recover the two halves of the 96-nm repeat, 48-nm particles were classified with a mask covering external complexes bound to protofilaments A02-A04. (b) Fitting the two halves of the 96-nm repeat into the subtomogram average from pig sperm shows that the bases of external axonemal complexes (like the nexin/dynein regulatory complexes and radial spokes) are resolved. (c) Maps of the two halves of the 96-nm repeat with corresponding FSC curves.

**Fig. S3.**
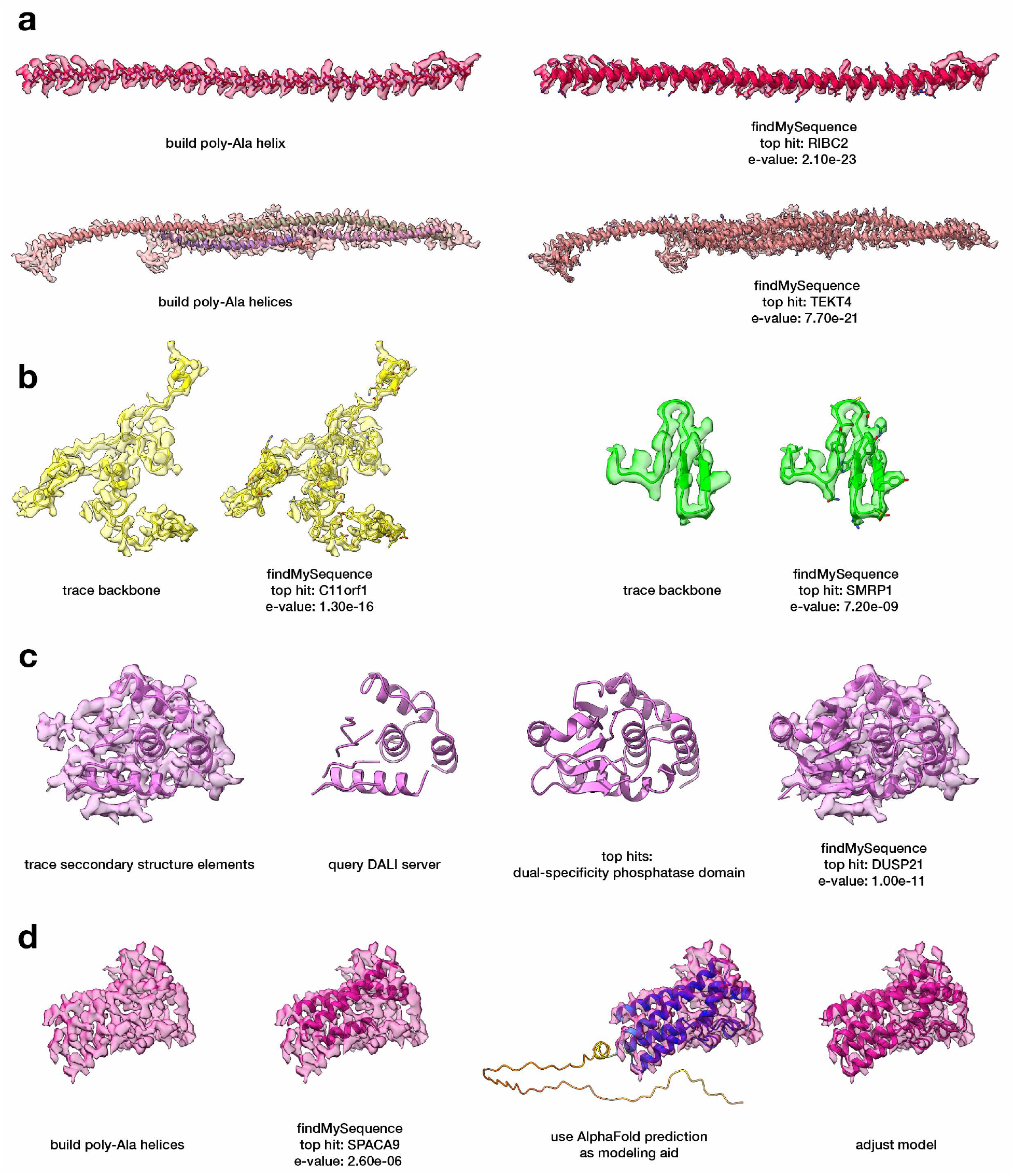
Examples of microtubule inner protein identification strategies. To identify most MIPs, poly-Ala models were manually built into the map, starting with secondary structure elements like helices or sheets as anchors and extending when the density permitted. The findMySequence program ^23^ was then used to query a database consisting of the 1500 most abundant proteins found by proteomics in our bovine sperm samples. **(a)** The workflow was first internally validated on known MIPs, like RIBC2 and tektins. **(b)** In most cases, backbone traces and findMySequence identified MIPs of varying lengths and folds. **(c)** In the case of DUSP21, running findMySequence directly on traced secondary structure elements did not yield a hit. Querying the DALI server ^64^ returned dual-specificity phosphatase domains as the top hits. The top hit (PDB 5Y16) ^65^ was mutated to polyAla, then fitted and refined into the density map of the unknown MIP. Running findMySequence on this model returned DUSP21 as a confident hit. **(d)** In the case of SPACA9, findMySequence could assign protein identity, but it was initially difficult to fully trace the backbone. Fortunately, the AlphaFold2 prediction ^66^ for SPACA9 was high-quality and could be used to facilitate model building.

**Fig. S4.**
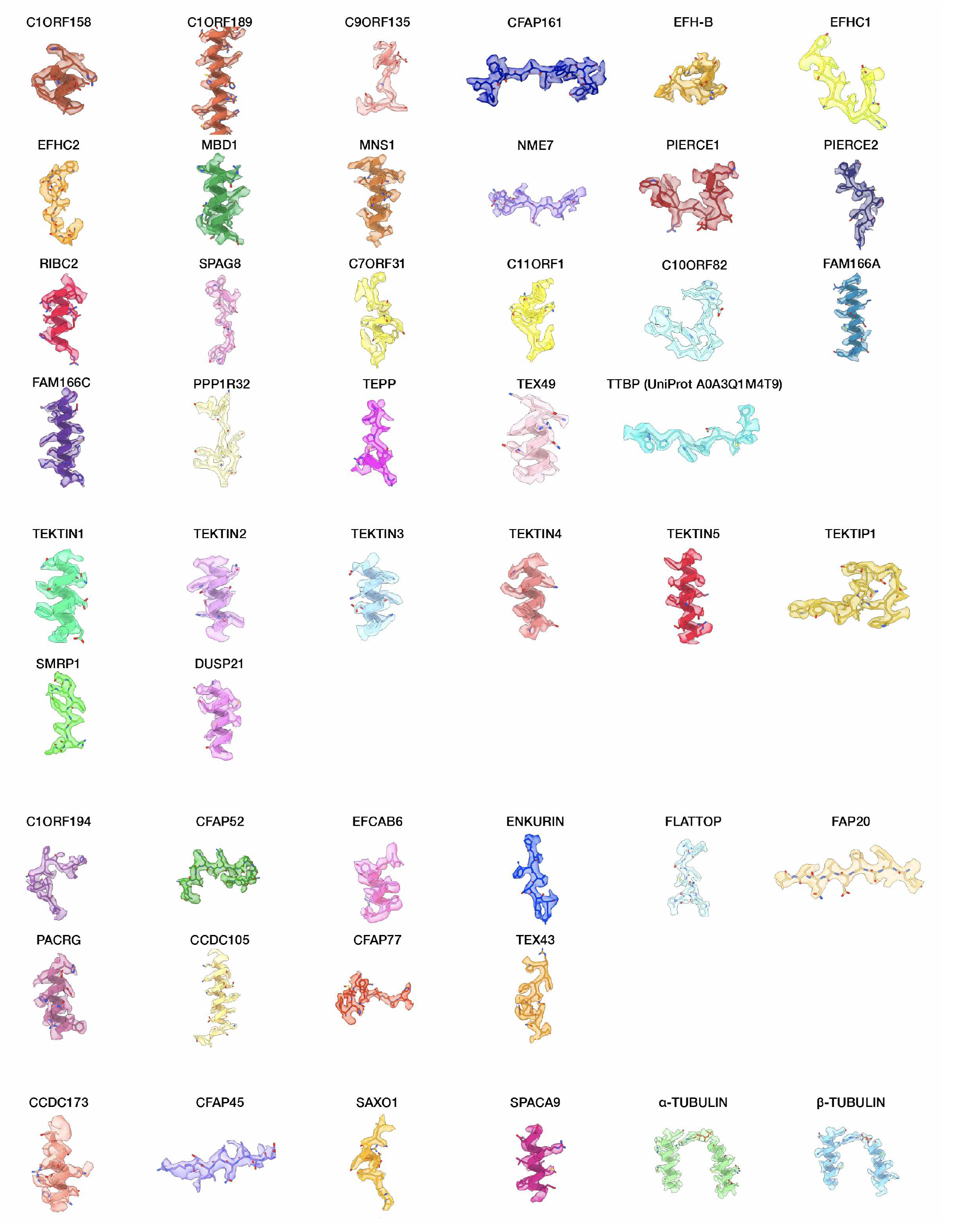
Examples of map quality for identified proteins in the bovine sperm DMT. Proteins are color-coded consistently throughout the manuscript.

**Fig. S5.**
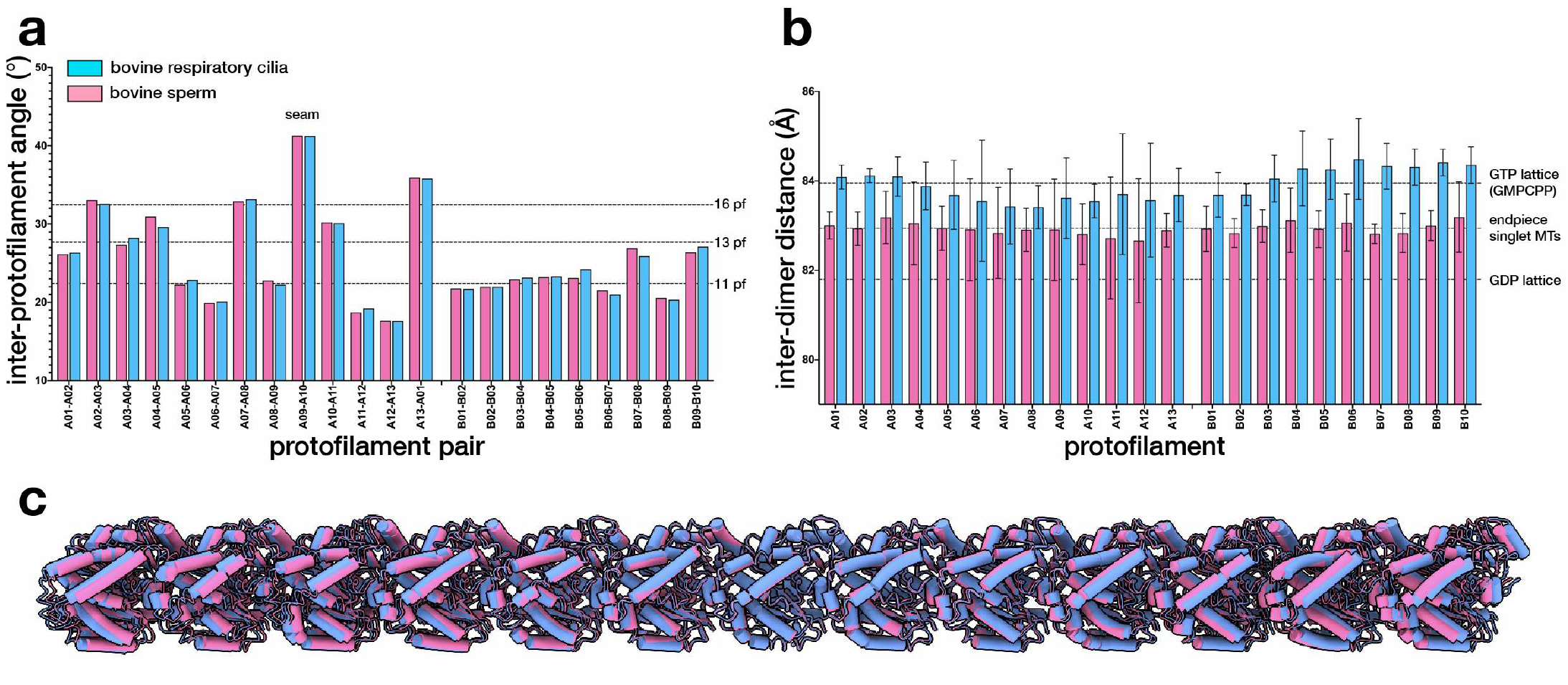
Comparing tubulin lattices between bovine tracheal DMTs and bovine sperm DMTs. **(a)** The inter-protofilament rotation angles are similar between DMTs from respiratory cilia (blue) and those from sperm (pink). **(b-c)** Bovine sperm DMTs have a more compact tubulin lattice and shorter inter-dimer distances than bovine tracheal DMTs.

**Fig. S6.**
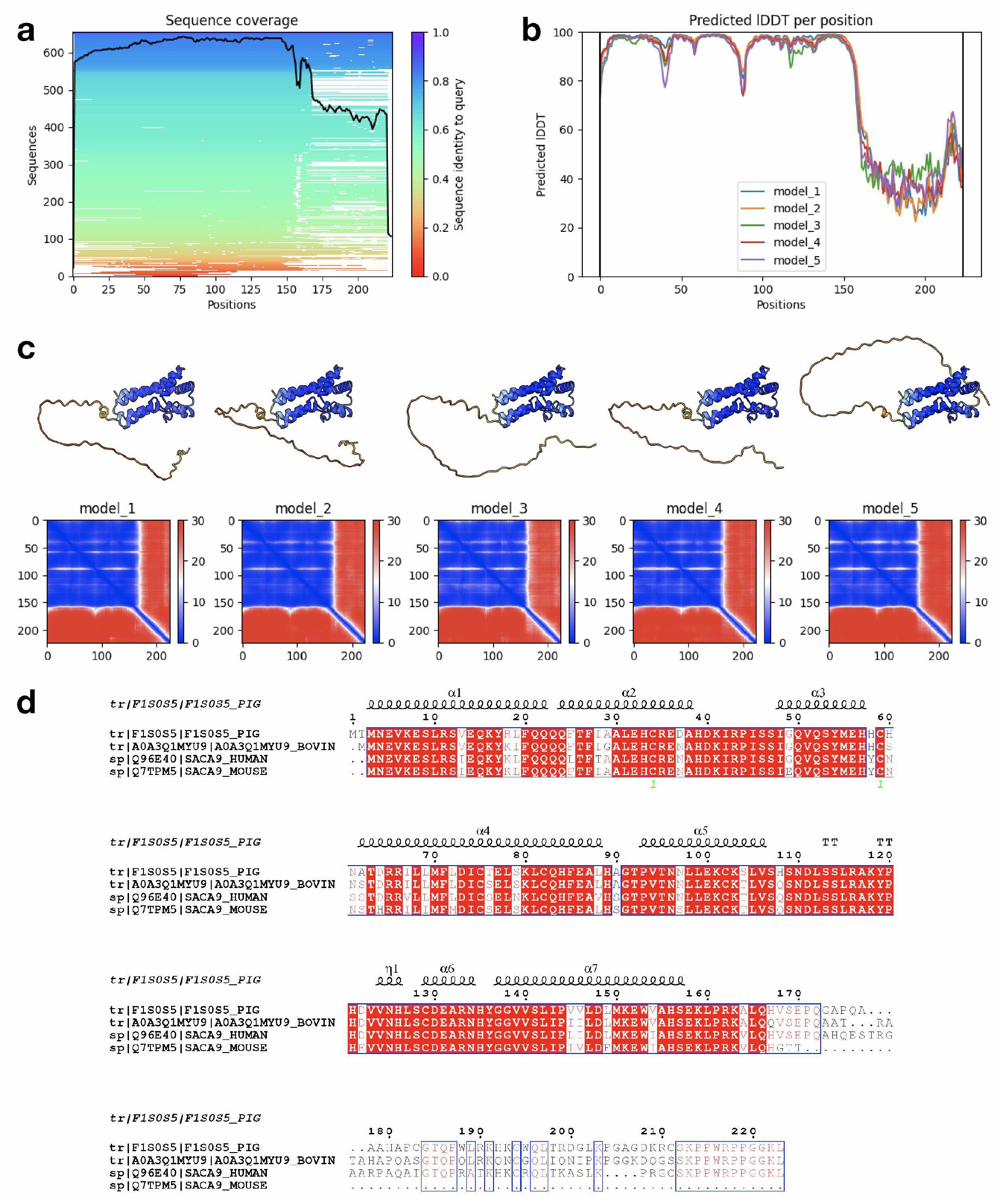
AlphaFold predictions and sequence conservation of SPACA9. **(a)** Sequence coverage, **(b)** predicted LDDT (pLDDT), **(c)** three-dimensional models colored according to pLDDT (top panels), and predicted aligned error (bottom panels) for the top five AlphaFold models of bovine SPACA9. Note how the N-terminal alpha-helical bundle is predicted with very high confidence. **(d)** Sequence alignment of SPACA9 from four mammalian species illustrating high conservation of the N-terminal domain and divergence of the C-terminal sequences.

**Fig. S7.**
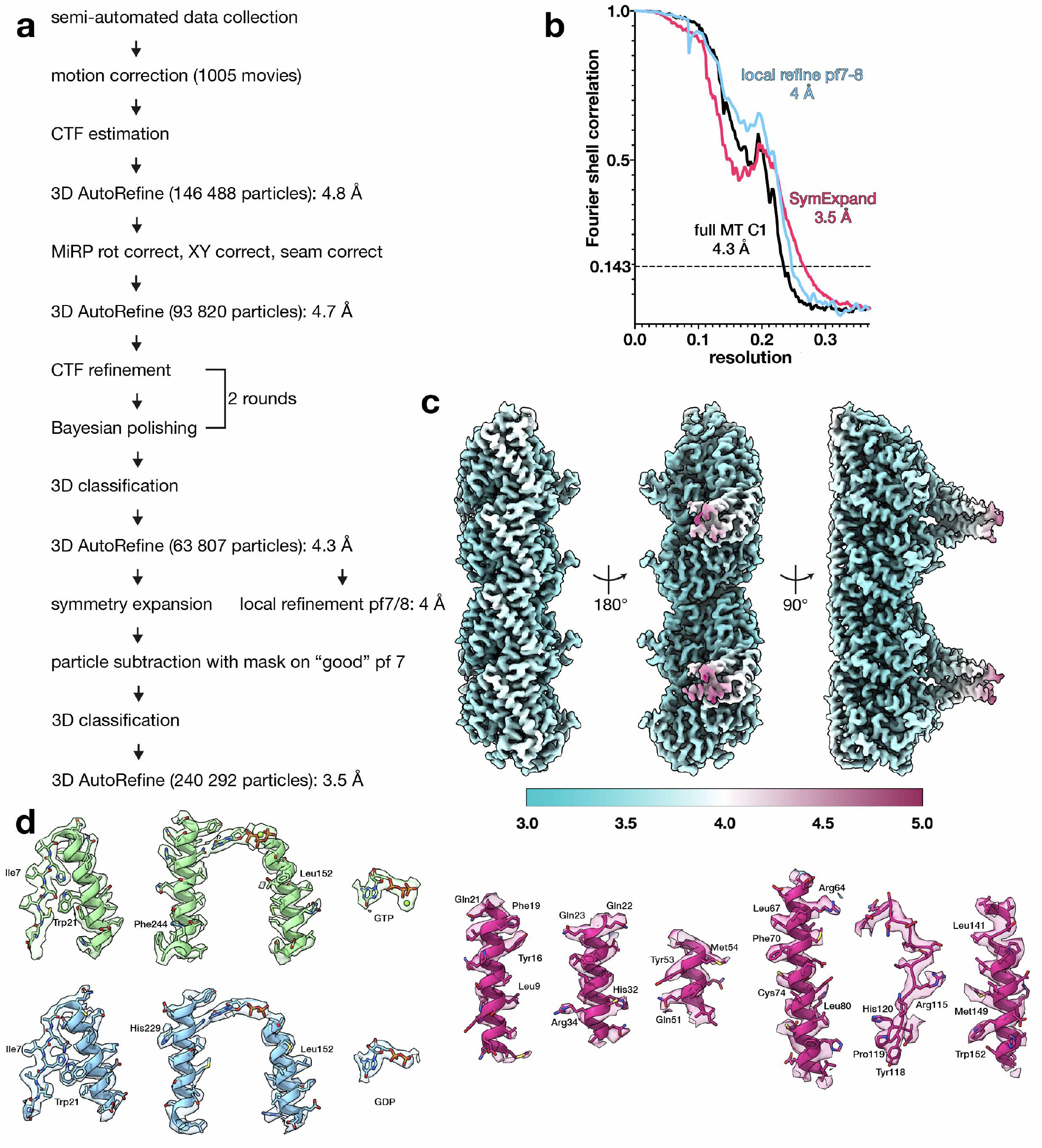
Cryo-EM processing workflow and map quality for endpiece singlet microtubules. **(a)** The cryo-EM workflow involved standard Relion refinements for a C1 helix, using a 20-Å subtomogram average of pig endpiece microtubules as a reference. After initial alignments, rotation angle correction, X/Y shift correction, and seam checking were performed according to the Microtubule Relion-based Pipeline (MiRP). Particles were then CTF-refined and Bayesian-polished. After 3D classification and selecting only the highest-resolution class, 3D refinement yielded a 4.3-Å C1 reconstruction of the full microtubule. Subsequent local refinement of protofilament 7/8 yielded at 4-Å map. After symmetry expansion and particle subtraction, followed by 3D classification to remove poorly-decorated particles, a refinement run focused on two tubulin dimers resulted in a 3.5-Å reconstruction. **(b)** Fourier shell correlation (FSC) plots of the full C1 reconstruction (black curve), the local refinement of protofilaments 7/8 (blue curve), and the protofilament reconstruction after symmetry expansion (pink curve). **(c)** Local resolution of the symmetry-expanded reconstruction, estimated by the “Local Resolution” job in Relion. **(d)** Examples of map quality in the symmetry-expanded map used for MIP identification and model building.

**Table S1.** Cryo-EM data collection, refinement, and validation statistics.

|  | bovine sperm doublet microtubules,<br>48-nm repeat<br>(EMDB-xxxx)<br>(PDB xxxx) | bovine sperm endpiece<br>singlet microtubules, 8-nm repeat<br>(EMDB-xxxx)<br>(PDB xxxx) |
| --- | --- | --- |
| **Data collection and processing** |  |  |
| Magnification | 130 000 X | 100 000 X |
| Voltage (kV) | 200 (dataset 1-4)<br>300 (dataset 5) | 200 |
| Electron exposure (e-/Å<sup>2</sup>) | ~48-50 | ~48-50 |
| Defocus range (µm) | -0.5 to -2.5 | -0.5 to -2.5 |
| Pixel size (Å) | 1.041 (dataset 1-4)<br>1.06 (dataset 5) | 1.359 |
| Symmetry imposed | C1 | C1, then helical symmetry-expanded |
| Initial particle images (no.) | 532 665 (8-nm) | 146 488 |
| Final particle images (no.) | 34 083 (48-nm) | 63 807 (C1), 240 292 (expanded) |
| Map resolution (Å) | ~3.7 | ~4.3 (C1), ~3.5 (expanded) |
| FSC threshold | 0.143 | 0.143 |
| **Refinement** |  |  |
| Initial model used (PDB code) | 7RRO, 7MIZ, AlphaFold, *de novo* | 3JAS, 7MIZ, AlphaFold |
| Model composition | 1 398 504 | 8139 |
| Non-hydrogen atoms | 175 378 | 1027 |
| Protein residues | GTP: 149, Mg: 149, | GTP: 1, Mg: 1, |
| Ligands | GDP: 153 | GDP: 1 |
| *B* factors (Å<sup>2</sup>) |  |  |
| Protein | 42.79 | 61.74 |
| Ligand | 45.41 | 45.86 |
| R.m.s. deviations |  |  |
| Bond lengths (Å) | 0.010 | 0.002 |
| Bond angles (°) | 0.945 | 0.548 |
| Validation |  |  |
| MolProbity score | 1.80 | 1.57 |
| Clashscore | 9.00 | 6.18 |
| Poor rotamers (%) | 0.59 | 0.00 |
| Ramachandran plot |  |  |
| Favored (%) | 95.36 | 96.45 |
| Allowed (%) | 4.58 | 3.45 |
| Disallowed (%) | 0.05 | 0.10 |

**Table S2. Top 1500 most abundant proteins identified in bovine sperm.** In the attached Excel file, Tab 1 reports the top 1500 most abundant proteins identified in bovine sperm, ordered according to iBAQ value. Highlighted rows indicate proteins identified as MIPs in the cryo-EM maps. Tab 2 reports the full bovine sperm proteome.

**Table S3.**
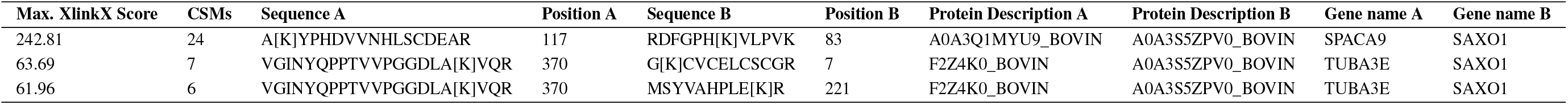
Cross-links identified between tubulin and SAXO1, and between SAXO1 and SPACA9.

**Movie S1.**
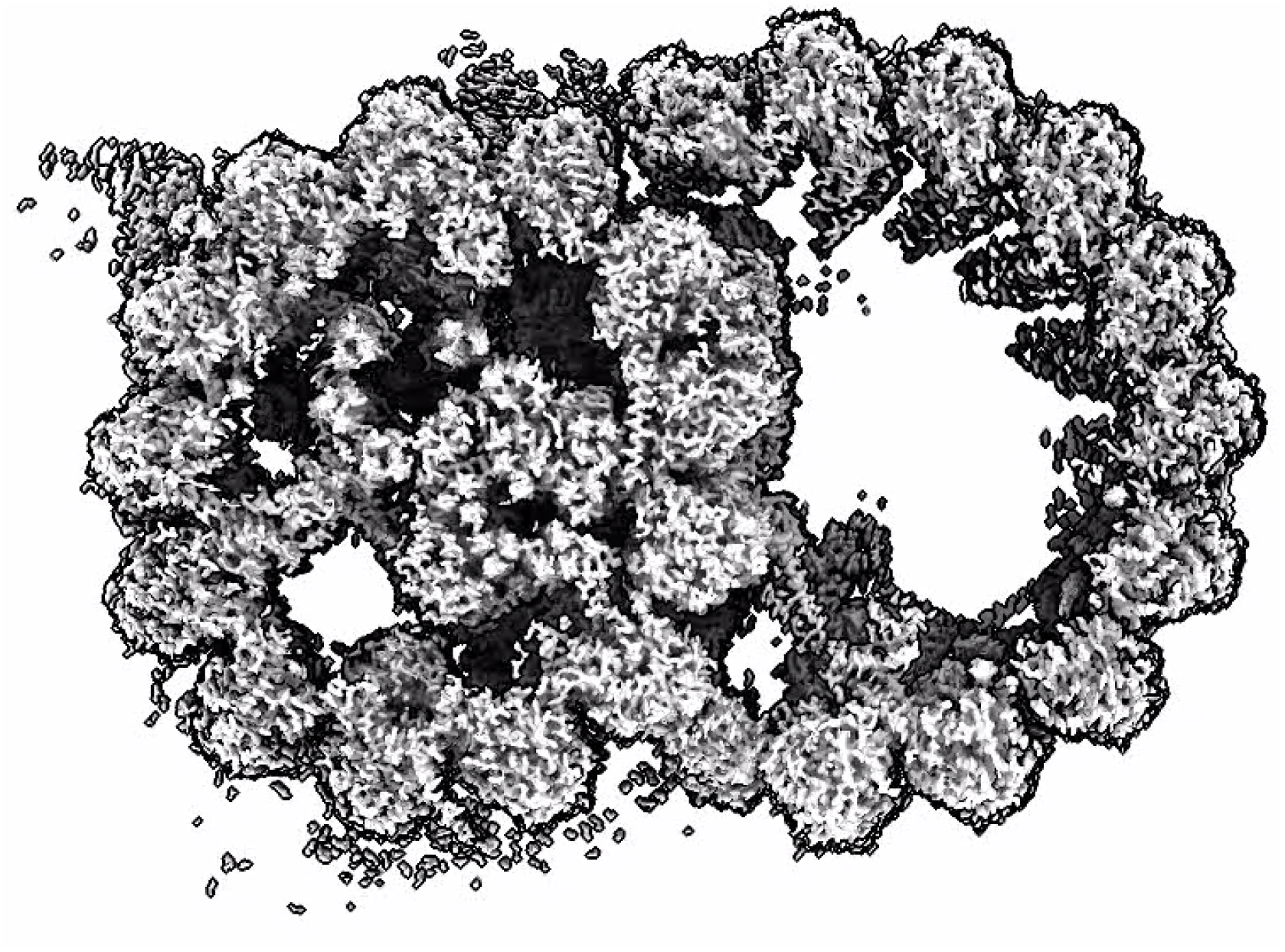
Structure of the 48-nm repeat of native axonemal doublet microtubules from mammalian sperm.

